# Parasite co-opts a ubiquitin receptor to induce a plethora of developmental changes

**DOI:** 10.1101/2021.02.15.430920

**Authors:** Weijie Huang, Allyson M. MacLean, Akiko Sugio, Abbas Maqbool, Marco Busscher, Shu-Ting Cho, Sophien Kamoun, Chih-Horng Kuo, Richard G.H. Immink, Saskia A. Hogenhout

## Abstract

Obligate parasites can induce complex and substantial phenotypic changes in their hosts in ways that favour their transmission to other trophic levels. However, mechanisms underlying these changes remain largely unknown. Here, we demonstrate how SAP05 protein effectors from insect-vectored plant pathogenic phytoplasmas take control of several plant developmental processes to simultaneously prolong host lifespan and induce witch’s broom-like proliferations of leaf and sterile shoots, organs colonized by phytoplasmas and vectors. SAP05 acts by mediating the concurrent degradation of SPL and GATA developmental regulators via a process that uniquely relies on hijacking the plant ubiquitin receptor RPN10 independently of substrate lysine ubiquitination. RPN10 is highly conserved among eukaryotes, but SAP05 does not bind insect vector RPN10. A two-amino-acid substitution within plant RPN10 generates a functional variant that is resistant to SAP05 activities. Therefore, one effector protein enables obligate parasitic phytoplasmas to induce a plethora of developmental phenotypes in their hosts.

## Introduction

Parasites are known to modulate specific processes in hosts to promote colonization and virulence. Whereas most pathogens colonize one host, a substantial number of obligate parasites require multiple hosts to complete their life cycles. These parasites often depend on their hosts feeding on each other, and fascinatingly, appear to have evolved mechanisms to induce developmental and behavioural modifications in their hosts that increase the chance of interactions among host trophic levels (Hughes and Libersat, 2019; Le Fevre et al., 2015). For example, the trematode, *Ribeiroia ondatrae*, causes severe limb abnormalities in Pacific treefrogs, such as the induction of growth of extra limbs or aborting limbs, thereby impairing movement of the frogs and increasing the risk of their predation by birds, which are the definitive hosts of the trematode parasite (Johnson et al., 1999). These parasites are spectacular examples of how the reach of genes can extend beyond the organisms to impact surrounding environments, a phenomenon known as the extended phenotype (Dawkins, 1982). However, the molecular mechanisms underpinning these parasite-enforced host modifications are largely unknown and there are ongoing debates about the extent to which these phenotypes are adaptive (Herbison et al., 2018; Johnson and Koshy, 2020).

One group of plant pathogens that are notorious for reprogramming their host development are members of *Candidatus* (Ca.) Phytoplasma (Al-Subhi et al., 2020; Doi et al., 1967; IRPCM, 2004; Lee et al., 2000; Sugio et al., 2011b), which comprises a diverse genus of bacteria that cause global socioeconomically important insect-transmitted diseases (EPPO, 2021). Phytoplasmas infect most vascular plant species and often induce massive changes in plant architecture, such as excessive proliferations of shoots and branches (witches’ brooms) and retrograde development of flowers into leaf-like organs (phyllody) (Hoshi et al., 2009; MacLean et al., 2011; Maejima et al., 2014; Sugio et al., 2011a). Plants exhibiting exhibit extensive architectural changes as the result of pathogen infections are described as ‘Zombies’, as the plants stop reproducing themselves and serve only as a habitat for the pathogens and their insect vectors (Al-Subhi et al., 2020; Cano et al., 2013; Du Toit, 2014; MacLean et al., 2014; Orlovskis and Hogenhout, 2016; Pecher et al., 2019; Rumpler et al., 2015; Sugio et al., 2011a). The three-way interactions among phytoplasmas, plants and insects, provide an excellent system to study the genetic basis of extended phenotypes created by obligate multi-host parasites (Huang et al., 2020; Sugio et al., 2011b).

Phytoplasmas and other vector-borne plant pathogens are strict obligates (Huang et al., 2020). Phytoplasmas have a dual host cycle that alternates between plants (Kingdom Plantae) and insects (Kingdom Animalia), and are reliant on both for survival (Hogenhout et al., 2008). In plants, phytoplasmas colonize the cytoplasm of vascular phloem sieve cells that transport nutrients to growing plant tissues (Doi et al., 1967). The bacteria spread systemically in plants via migration through the phloem cell sieve pores (Musetti et al., 2013). Sap-feeding insects that feed from the phloem, predominantly leafhoppers, planthoppers and psyllids of the order Hemiptera, are efficient phytoplasma vectors (Weintraub and Beanland, 2006). In these insects, phytoplasmas colonise most tissues, including the salivary glands through which the bacteria are introduced into the plant vasculature during phloem feeding (Frost et al., 2011; Koinuma et al., 2020). Whereas phytoplasmas can be deleterious to their insect vectors, the phytoplasmas often have beneficial effects on their vectors, especially in established pathosystems where the bacteria and the insects have co-evolved over long periods of time (Beanland et al., 2000; Malembic-Maher et al., 2020; Nault, 1990).

Progress in the characterization of phytoplasma virulence factors greatly accelerated by the ability of some phytoplasmas to colonise the model plant *Arabidopsis thaliana*. One of these phytoplasmas is Aster Yellows phytoplasma (AYP) strain Witches’ Broom (AY-WB; *Ca*. Phytoplasma asteris) (Hogenhout et al., 2008; Sugio et al., 2011b; Zhang et al., 2004). The main vector of AYPs in North America is the polyphagous aster leafhopper *Macrosteles quadrilineatus*, which migrates over long distances and transmits the bacteria to various crops (Frost et al., 2013a, b). AYPs and *M. quadrilineatus* colonise a broad range of plant species, including oilseed rape and other members of the *Brassicaceae*, as well as lettuce, carrots and several cereals (CABI, 2021). Aster yellows phytoplasmas induce witches’ brooms and phyllody symptoms and their occurrence can be high, sometimes contributing to loss of entire crop productions (Frost et al., 2013a). Phytoplasma cause these symptoms by secreting proteins, known as effectors and unload them from the phloem to adjacent plant tissues, such as shoot and apical meristems, where cells are programmed to give rise to various plant organs (Arashida et al., 2008; Bai et al., 2009; Hoshi et al., 2009; Kakizawa et al., 2004; MacLean et al., 2011). Mining of the AY-WB genome for potential effectors resulted in the identification of 56 candidate effector genes, named secreted AY-WB proteins (SAPs) (Bai et al., 2009). To date only a few of these have been characterized. SAP11s bind and destabilize *A. thaliana* TCP transcription factors resulting in leaf shape changes and stems proliferations (Sugio et al., 2011; Sugio et al., 2014). SAP54 binds and degrades *A. thaliana* MADS-box transcription factors via co-opting proteasome RAD23 shuttle factors leading to the development of flowers that remain leafy and the indeterminate growth of stems from the flower-like organs (MacLean et al., 2011; MacLean et al., 2014). Homologs of these effectors have been found in divergent phytoplasmas and shown to degrade TCPs and MADS-box transcription factors of other plant species (Chang et al., 2018; Iwabuchi et al., 2020; Kitazawa et al., 2017; Lu et al., 2014; Maejima et al., 2014; Pecher et al., 2019; Tan et al., 2016; Wang et al., 2018). However, the phenotypes caused by SAP11, SAP54 and their homologs do not account for all the extensive developmental phenotypes caused by phytoplasmas, such as prolonged lifespan and witch’s broom type tissue proliferations other than stems.

Here, we discovered that a new phytoplasma effector SAP05 binds and mediates degradation of multiple members of the two distinct SPL and GATA transcription factor families, SPL and GATA, leading to delayed plant aging and the simultaneous proliferations of vegetative tissue and shoots. SAP05 varies among phytoplasmas. Some strains have two SAP05 homologs of which one bind SPLs and the other GATAs accounting for complex phenotypic outcomes. The SAP05 effectors mediate degradation through a unique lysine ubiquitination-independent mechanism, by co-opting the 26S ubiquitin receptor RPN10 that is highly conserved across eukaryotes. Remarkably, SAP05 does not bind the RPN10 of insect phytoplasma insect vectors and only two RPN10 amino acids define binding specificity. These two amino acids are one of few lowly conserved sequences between plant and animal RPN10. We used this information to engineer a functional variant of plant RPN10 that has lower affinity for SAP05 and that confers plant resistance to SAP05 activity during phytoplasma infection. This work shows that one phytoplasma effector co-opts one proteasome protein to degrade multiple developmental regulators and to reprogram plant development. This induces a plethora of adaptive phenotypic changes in their plant hosts.

## Results

### SAP05 alters plant architecture and reproduction

As part of functional screens with candidate AY-WB phytoplasma effectors, we found that SAP05 perturbs plant developmental processes by analysing phenotypes of stable transgenic *A. thaliana* lines that constitutively express the *SAP05* gene corresponding to the mature part of the protein (without signal peptide) under control of the constitutive Cauliflower mosaic virus (CaMV) *35S* promoter. These *SAP05*-expressing plants exhibited a range of architectural differences compared to control plants that express the gene for green fluorescent protein (GFP) from the same promoter (Figures 1 and S1). During vegetative growth, SAP05 plants displayed accelerated leaf initiations and produced more rosette leaves (Figures 1A, 1E, S1A and S1C). In addition, the leaves of mature rosettes lacked serrated edges that are present in wild-type and *GFP*-expressing plants (Figures 1A and S1A). A closer examination revealed that the appearance of abaxial trichomes, which is characteristic of adult leaves, was also delayed in SAP05 plants (Figures 1E and S1C). However, SAP05 plants initiated budding and flowering no later than the control plants, under both short-day and long-day conditions (Figures 1F and S1D). SAP05 plants produced more lateral shoots and secondary branches, and these plants were reduced in height (Figures 1C, 1D, 1G and 1H). At 12 weeks in LD, GFP plants started to senesce, whereas the SAP05 plants continued to grow (Figure 1D), suggesting that SAP05 delays plant senescence. Twenty-six of 32 independently transformed *p35S::SAP05* lines developed abnormal flowers and were greatly compromised in fertility (Figure 1C, inset), with extremely bushy plants showing complete sterility (Figure S1E).

**Figure 1.**
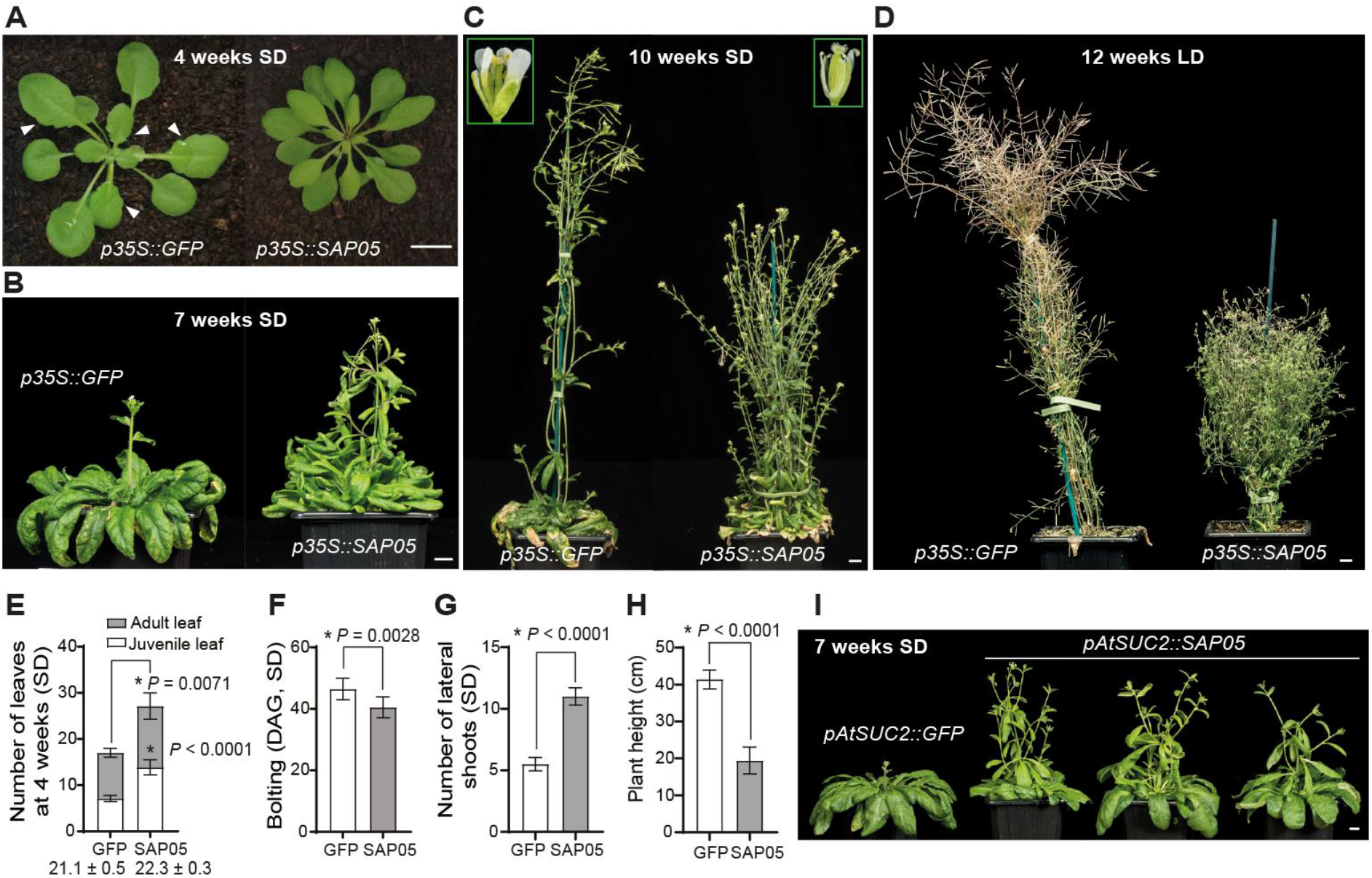
Phytoplasma effector SAP05 induces excessive shoot proliferation and sterility in *A. thaliana*. (A-D) Representative images of transgenic plants expressing SAP05 or GFP (control). Plants were grown under short-day (SD) or long-day (LD) conditions, and images were obtained at 4 weeks (A), 7 weeks (B),10 weeks (C) and 12 weeks (D) after germination. Arrowheads in (A) indicate leaf serrations of GFP plants as opposed to the smoother leaf edges of SAP05 plants. Insets in (C) show enlarged images of mature flowers with a same magnification. Scale bars, 1 cm. For plants grown under long-day (LD) conditions, see also Figure S1. (E-H) Statistical analysis of phenotypes shown in (A-C): numbers of rosette leaves of 4-week-old plants (E) and time of shoot emergence from rosettes (bolting time; F), number of shoots emerging from rosettes (lateral shoot number; G) and plant height (H) of 10-week-old SD plants. Numbers under the bars in (E) indicate the time when the first abaxial trichome appeared. DAG, day after germination. *, *P* < 0.05, two-tailed unpaired Student’s *t-*tests. (I) Morphology of a GFP control plant and plants from three independent *A. thaliana* transgenic lines expressing SAP05 under the control of the phloem-specific *AtSUC2* promoter. Images were obtained at 7 weeks after germination. Scale bar, 1 cm.

Because phytoplasmas secrete effector proteins into the phloem cells, we generated stable transgenic *A. thaliana* lines expressing SAP05 from the phloem-specific *AtSUC2* promoter (Mathieu et al., 2007). The *pAtSUC2::SAP05* plants exhibited similar architectural changes to the *p35S::SAP05* plants, exhibiting stunting, bushiness and sterility (Figure 1I), phenotypes that resemble the witch’s broom symptoms typically observed in phytoplasma-infected plants.

### SAP05 mediates plant transcription factors of the SPL and GATA families for destabilization via the ubiquitin receptor RPN10

To identify potential SAP05 targets in plants, a yeast two-hybrid (Y2H) screen against an *A. thaliana* seedling library was conducted. We found that SAP05 interacts with several *A. thaliana* zinc-finger transcription factors, specifically GATA and SPL transcription factors (Figures 2A-2C and Tables S1, S2). Of the 29 GATA and 16 SPL family members in *A. thaliana*, 26 GATAs and 12 SPLs were successfully cloned and shown to interact with SAP05 in Y2H assays, indicating that SAP05 most likely binds all members of both families. SPLs regulate plant developmental phase transitions, and most are developmentally regulated by microRNA156 (miR156) (Xu et al., 2016), whereas GATA proteins regulate photosynthetic processes, leaf development and flower organ development (Ding et al., 2015; Ranftl et al., 2016; Richter et al., 2010). The SPL and GATA transcription factor families evolved independently, though both have zinc-finger (ZnF) DNA-binding domains. The ZnF domain of the GATA family contains one C4-type zinc-binding site of approximately 50 residues and is conserved among plants, animals and fungi, whereas that of SPL proteins is approximately 80 amino acids, contains two zinc-binding sites of the C3H and the C2HC types and is specific to plants. The ZnF domains of SPLs and GATAs are both sufficient to mediate SAP05 binding in Y2H experiments (Figure 2C).

**Figure 2.**
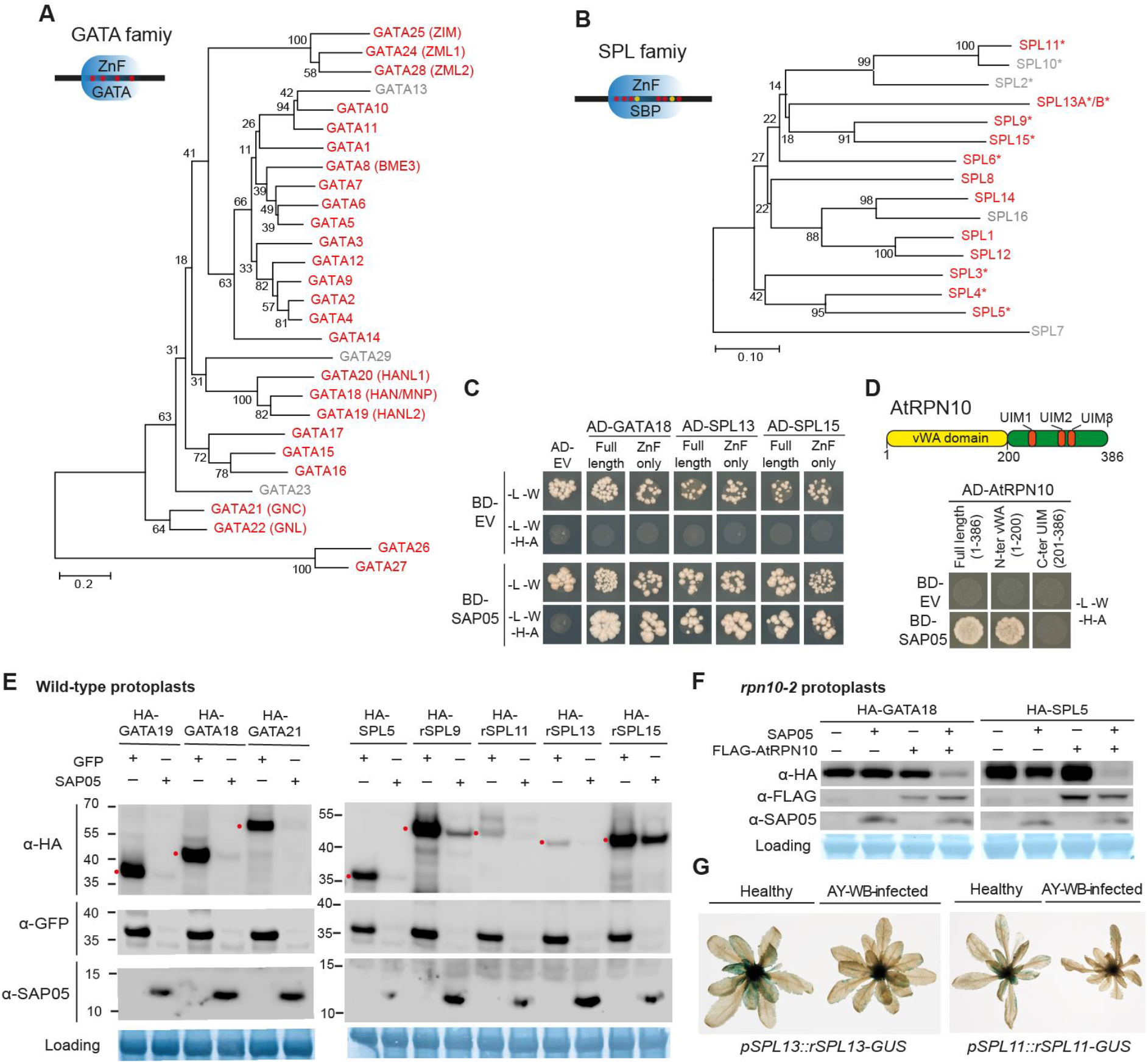
SAP05 destabilizes plant transcription factors of the SPL and GATA families via interaction with the ubiquitin receptor AtRPN10. (A and B) SAP05 interacts with most members of the *A. thaliana* GATA (A) and SPL (B) transcription factor families in Y2H assays. The phylogenies show SAP05 interactors in red and those that were not tested or had autoactivation activities in grey. Conserved zinc-finger (ZnF) domains are shown in the upper left corners with red and yellow dots indicating cysteine and histidine residues, respectively. SBP, SQUAMOSA promoter-binding protein. *Regulated by miR156. (C) SAP05 interacts with the ZnF domains of GATAs and SPLs in Y2H assays. EV, empty vector control. AD, GAL4-activation domain. BD, GAL4-DNA binding domain. Yeast growth on medium lacking leucine and tryptophan (-L-W) indicates presence of AD and BD constructs and on medium lacking leucine, tryptophan, histidine and alanine (-L-W-H-A) interactions between the AD and BD fusion proteins. (D) SAP05 interacts with AtRPN10 in Y2H assays. Upper panel, graphical representation of AtRPN10 domains with locations indicated in amino acids underneath. vWA, von Willebrand factor type A domain; UIM, ubiquitin-interacting motifs. See Fig. S10A for yeast growth on -L-W medium. (E) Western blot analysis of SAP05-mediated degradation of GATA and SPL proteins in protoplasts from wild type *A. thaliana*. GFP, control; HA, hemagglutinin tag; rSPL, miR156-resistant form. Numbers at left indicate molecular weight markers in kilodaltons. Red dots at left of the blots indicate the expected sizes of the transiently expressed proteins. Protein loading was visualized using Amido black staining. (F) Western blot analysis of SAP05 degradation assays in *rpn10-2* protoplasts. (G) GUS staining produced by the GUS-fusions of the miR156-resistant forms of SPL11 (rSPL11) and SPL13 (rSPL13) in the transgenic *A. thaliana* is reduced in AY-WB-infected plants at right compared to non-infected plants at left.

In addition, the *A. thaliana* 26S proteasome subunit RPN10 was identified as a potential SAP05 interactor in the Y2H screen (Figure 2D and Table S1). RPN10 is located within the 19S regulatory particle of the proteasome and serves as one of the main ubiquitin receptors recruiting ubiquitinated proteins for proteasomal degradation (Fu et al., 1998; van Nocker et al., 1996). Recently, RPN10 was also shown to mediate the autophagic degradation of inactive 26S proteasomes (Marshall et al., 2015). RPN10 has two main domains (Riedinger et al., 2010). The N-terminal vWA (von Willebrand factor type A) domain is required for RPN10 docking to the proteasome (Mayor et al., 2007). The C-terminal half with ubiquitin-interacting motifs (UIM) is involved in binding to ubiquitin chains or the autophagic factor ATG8 that directed either ubiquitinated substrates or inactive proteasomes, respectively, for degradation (Figure 2D). We found that SAP05 interacts with full-length protein and the vWA domain, but not the UIM domains of AtRPN10 (Figure 2D).

Given that SAP05 interacts with the 26S proteasome subunit RPN10, we investigated if SAP05 degrades GATA and SPL transcription factors in plant cells. To this end, *GATA* or *SPL* genes and *SAP05* (or *GFP* as control) were transiently co-expressed under control of the constitutive 35S promoter in *A. thaliana* protoplasts. GATA proteins were absent or less abundant in the presence of SAP05 compared to GFP (Figure 2E, left panel). Similarly, the abundance of five SPL protein was visibly reduced in the presence of SAP05 (Figure 2E, right panel). To investigate the role of AtRPN10 in the SAP05-mediated destabilization of plant transcription factors, we made use of the existing and well-described *A. thaliana* loss-of-function *rpn10* mutant line *rpn10-2* (Lin et al., 2011; Smalle et al., 2003). In the protoplasts generated from *rpn10-2* plants, SAP05 no longer degraded AtGATA18/HAN or AtSPL5, whereas when *AtRPN10* was reintroduced into the protoplasts, these plant transcription factors were degraded in the presence of SAP05 (Figure 2F). Therefore, AtRPN10 is required for the SAP05-mediated degradation of plant targets.

To investigate whether GATAs and SPLs are degraded during phytoplasma infection, we made use of lines that express miR156-resistant forms of *SPL11* and *SPL13* (r*SPL11* and r*SPl13*, respectively) translationally fused with a β-glucuronidase protein (GUS) under the control of their native promoters (Xu et al., 2016). In line with previous findings, *rSPL11::GUS* and *rSPL13::GUS* were expressed in young leaves but not in fully expanded leaves. Newly emerged or developing leaves of phytoplasma-infected plants had visibly reduced GUS activities compared to un-infected plants (Figure 2G), whereas the expression levels of those two genes in developing leaves did not differ between healthy and diseased plants (Figure S2). These data further support that the SPL transcription factor proteins are degraded during phytoplasma infection in plants.

### SAP05 bridges host transcription factors to RPN10 for degradation via the 26S proteasome

Next, we investigated how SAP05 interaction with AtRPN10 leads to the degradation of SPLs and GATAs. Addition of MG132 and Bortezomib, two potent proteasome inhibitors, inhibited the SAP05-mediated destabilization of representative GATA and SPL proteins tested (Figures 3A-B). In contrast, two autophagy inhibitors, 3-Methyladenine and E-64d, did not interfere with the SAP05-mediated degradation (Figures 3A-B). Therefore, in the presence of SAP05, GATA and SPL proteins are targeted for destabilization in the host 26S proteasome.

**Figure 3.**
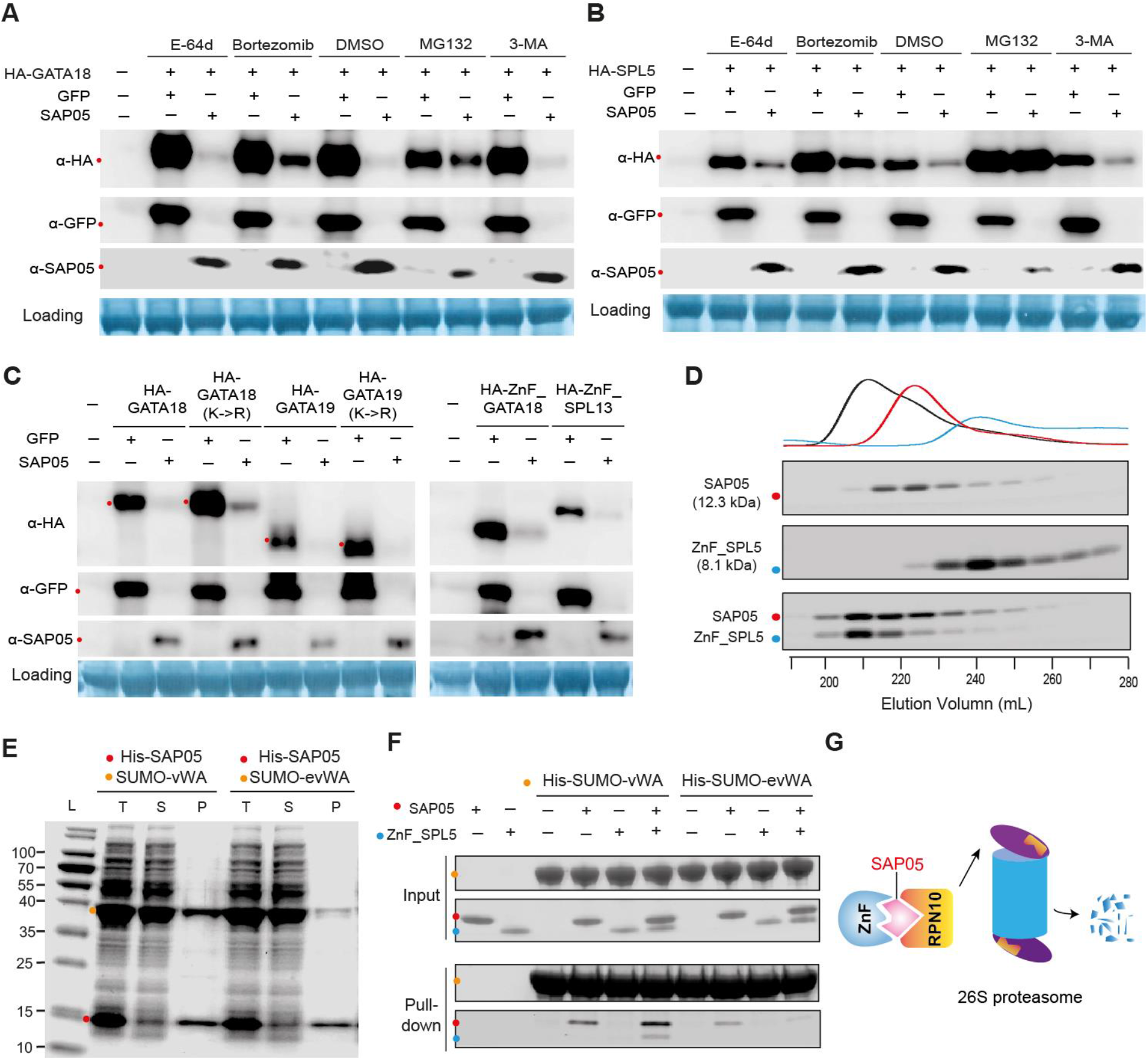
SAP05 hijacks the host ubiquitin receptor RPN10 to destabilize plant GATA and SPL transcription factors in the 26S proteasome. (A and B) 26S proteasome inhibitors reduce SAP05-mediated degradation of plant GATA and SPL in *Arabidopsis* protoplasts. MG132 and Bortezomib, 26S proteasome inhibitors. 3-Methyladenine, (3-MA) and E-64d, autophagy inhibitors. DMSO, control without inhibitors. (C). SAP05-mediated degradation does not require lysines in targeted proteins and only requires the SAP05-binding ZnF domain on targets. K->R, all lysines replaced by arginines. (D) Direct interaction of SAP05 and the ZnF domain of AtSPL5 in gel filtration assays. Coomassie stained SDS-PAGE gels with eluted fractions from gel filtration columns are shown. The top graph displays protein elution profiles (UV absorbance at 280 nm) from the columns over time. Coloured dots at left of the gels indicate the expected sizes of recombinant proteins in the gels. (E) IMAC co-purification of His-tagged SAP05 and the vWA domain of AtRPN10. evWA: vWA domain mutant with reduced affinity to SAP05 in Y2H assays (see Figure 5B). L: ladder. T: total cell extract. S: soluble fraction. P: protein purified by IMAC. Numbers at left indicate molecular weight markers in kilodaltons. (F) His-tag pull-down of ternary complexes of AtRPN10, SAP05 and the AtSPL5 ZnF domain. His-SUMO-tagged vWA domain or evWA domain were used as bait in IMAC to pull-down untagged SAP05 and/or the ZnF domain of AtSPL5. The coloured dots at left indicate the expected locations of recombinant proteins included in the assays on the blots. (G). A schematic overview of how SAP05 interaction with both ZnF and RPN10 leads to degradation of SPL and GATA transcription factors *in planta*. Arrow at left illustrates RPN10 interaction with the 19S regulatory particle of the plant 26S proteasome and arrow at right degradation of plant SPLs and GATAs by proteasome.

Given that RPN10 function as one of the main ubiquitin receptors, recruiting poly-ubiquitinated proteins for proteasomal degradation (Fu et al., 1998; Lin et al., 2011), we investigated whether the ubiquitination of lysine residues within SAP05 targets is necessary for their degradation. AtGATA18 and AtGATA19 proteins in which all lysines were replaced by arginines were also degraded in an SAP05-dependent manner (Figure 3C, left panel). Therefore, lysine ubiquitination of substrates is not required for SAP05-mediated degradation.

In addition, SPL or GATA ZnF domains alone that are targeted by SAP05 were also degraded in the presence of SAP05 (Figure 3C, right panel), suggesting that direct protein-protein interaction is key to substrate degradation initiation. To test whether there is direct binding between SAP05 and its plant targets, recombinant proteins or specific domains were expressed in *Escherichia coli* for detecting protein-protein interaction. Firstly, the interaction between SAP05 and the ZnF domain of AtSPL5 was confirmed in gel filtration assays. The two protein co-eluted in gel filtration with an elution profile distinct from those of the two proteins alone (Figure 3D), suggesting of stable complex formation *in vitro*. Moreover, a SUMO-tagged vWA domain was co-purified with His-tagged SAP05 from *E. coli* using immobilized metal affinity chromatography (IMAC) for the affinity purification of His-tagged fusion proteins (Figure 3E). In contrast, a mutated vWA domain (38GA39->HS, evWA) that does not interact with SAP05 in Y2H assays (further discussed below, see Figure 5B), was less enriched during co-purification (Figure 3E). Therefore, SAP05 also binds to the vWA domain *in vitro*. Finally, His-SUMO-tagged vWA domain pulled down the ZnF domain of AtSPL5 in the presence of SAP05, but not in the absence of this phytoplasma effector, and the evWA domain pulled down less SAP05 and did not pull down the ZnF domain in the presence of SAP05 (Figure 3F). Therefore, SAP05 forms bridge between the AtSPL5 ZnF and AtRPN10 vWA to generate a ternary complex. These results also indicate that SAP05 directly targets proteins that it interacts with for degradation and that lysine ubiquitination is not required for this degradation. Indeed, SAP05 fused to a GFP-nanobody (Rothbauer et al., 2006), a single-chain antibody domain that specifically recognizes GFP, also degraded GFP in *A. thaliana* protoplasts (Figure S3).

Taken together, these results indicate that SAP05 mediates SPL/GATA degradation in a lysine ubiquitination-independent manner by hijacking the plant host 26S proteasome component RPN10 (Figure 3G).

### Concurrent destabilization of SPLs and GATAs by SAP05 effectors decouples plant developmental transitions

To better understand how SAP05 has evolved to concurrently target two unrelated transcription factor families, we mined available genome sequences of divergent phytoplasmas for *SAP05* genes. *SAP05* homologs were identified in 17 phytoplasmas (Figure 4A). These phytoplasmas belong to the three major clades and 7 out of 10 16S rDNA (16Sr) groups of the phytoplasma phylogeny. SAP05 homologs were more frequently found among phytoplasmas compared to SAP54 and SAP11 homologs (10 and 15 phytoplasmas, respectively) (Figure 4A). The majority of phytoplasmas have one SAP05 homolog and 6 phytoplasmas have two SAP05 homologs. Most of these homologs interacted with both SPLs and GATAs (Figure 4B). However, for witches’ broom disease of lime (WBDL) phytoplasma and peanut witches’-broom (PnWB) phytoplasmas that both have two SAP05 genes, one SAP05 interacted only with SPLs and the other only with GATAs. Moreover, SAP05 of *Ca*. Phytoplasma mali (AT) only interacted with SPLs. Consistent with these binding specificities in yeast, the SAP05 homologs degraded one or both representative members of the SPL and GATA families in protoplast destabilization assays (Figure 4C). Phylogenetic analyses of the SAP05 effectors revealed distinct subclades of SAP05 homologs that bind and degrade both transcription factors or only SPLs or GATAs (Figure 4D). Together these data indicate that some SAP05 homologs have evolved to differentially interact and degrade plant SPL and GATA transcription factors.

**Figure 4.**
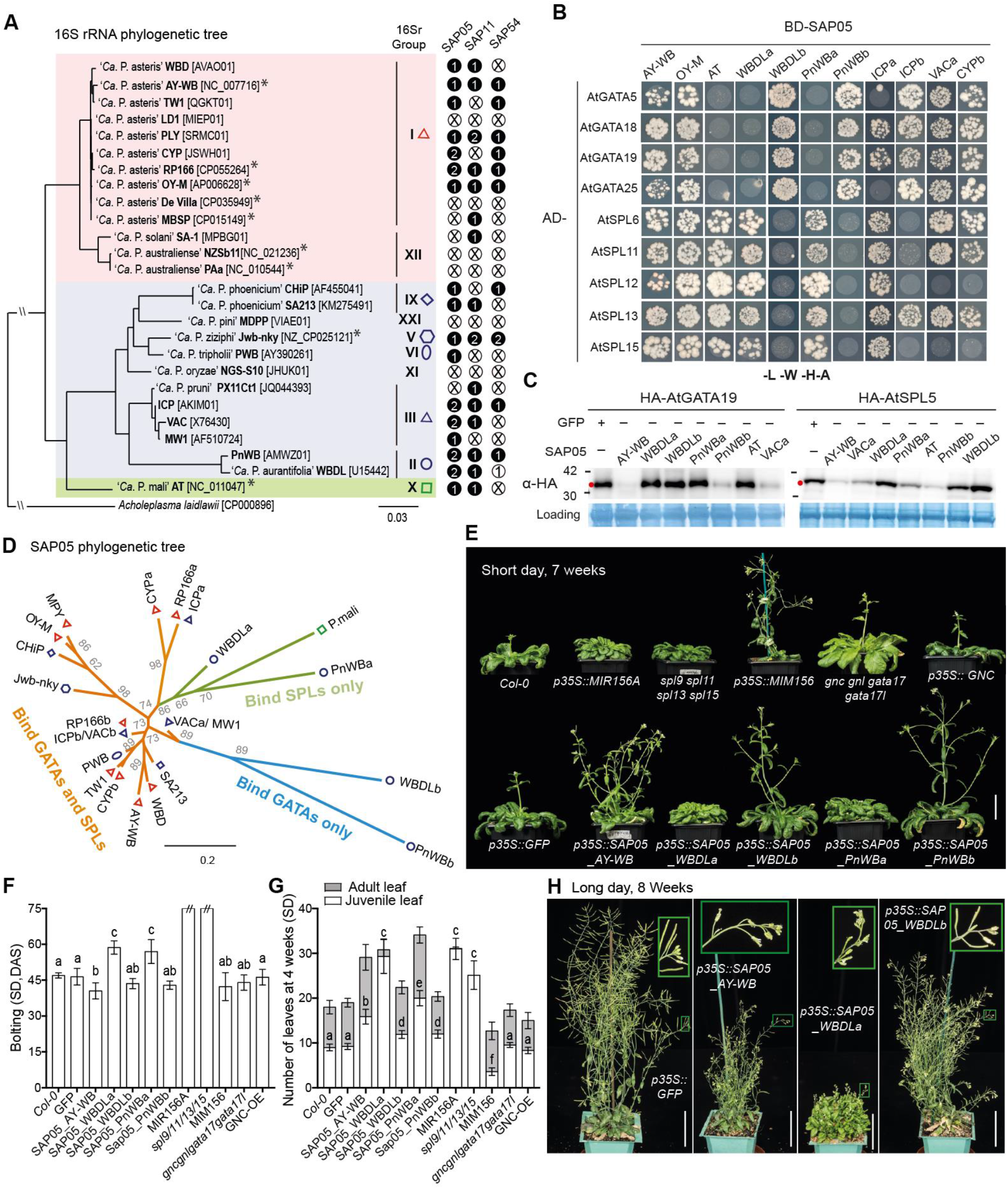
Phytoplasma SAP05 family effectors differentially bind and degrade plant SPL and GATA transcription factors. (A) SAP05 homologs are present in divergent phytoplasmas. The phylogenetic tree is based on the phytoplasma 16S rRNA gene alignment (Chung et al., 2013). The three distinct clades (indicated in pink, purple and green) and 16Sr groups are shown. The black circles indicate presence and the number of SAP05, SAP11 and SAP54 effector genes found in each phytoplasma and the white circles with crosses absence of these genes. *Phytoplasmas for which genomes were sequenced to completion. Shapes in different colours indicate phytoplasma subgroups with the presence of SAP05 genes. (B) Interactions of SAP05 homologs from divergent phytoplasmas and representative *A. thaliana* (At) GATA and SPL proteins in Y2H assays. See legend of Fig. 2 for abbreviations. Yeast growth on -L-W medium was shown in Fig. S10B. (C) SAP05 homologs of divergent phytoplasmas degrade *A. thaliana* (At) GATA19 and SPL5 in *A. thaliana* protoplasts. See legend of Fig. 2 for abbreviations. (D) SAP05 phylogenetic tree based on alignment of SAP05 amino acid sequences. Phytoplasma names and symbols to indicate 16Sr groups are shown in Fig. 4A. Branch lengths correspond to number of amino acid changes. SAP05 on orange branches bind both SPLs and GATAs in Y2H assays, and those on green and blue branches only GATAs or SPLs, respectively. Boot strap values are indicated at the nodes. (E) Morphologies of wildtype plants, *GFP*-expressing control plants, plants overexpressing different SAP05 homologs and transgenic plants with altered expression of *MIR156/SPL* or *GATA* genes. Plants were grown under short-day conditions. Scale bar, 4 cm. (F and G) Statistical analysis of phenotypes shown in (E): numbers of rosette leaves of 4-week-old plants (F) and time of shoot emergence from rosettes (bolting time; G). ‘//’ indicate that the plants did not bolt at the time of observation. Data are mean ± s.d.; different letters indicate significant difference based on multiple comparisons (Turkey method) after ANOVA. (H) Morphologies of GFP control plants and plants expressing different SAP05 homologs grown under long day conditions. Insets show enlraged images of inflorescence on mature plants. Scale bar, 4 cm.

**Figure 5.**
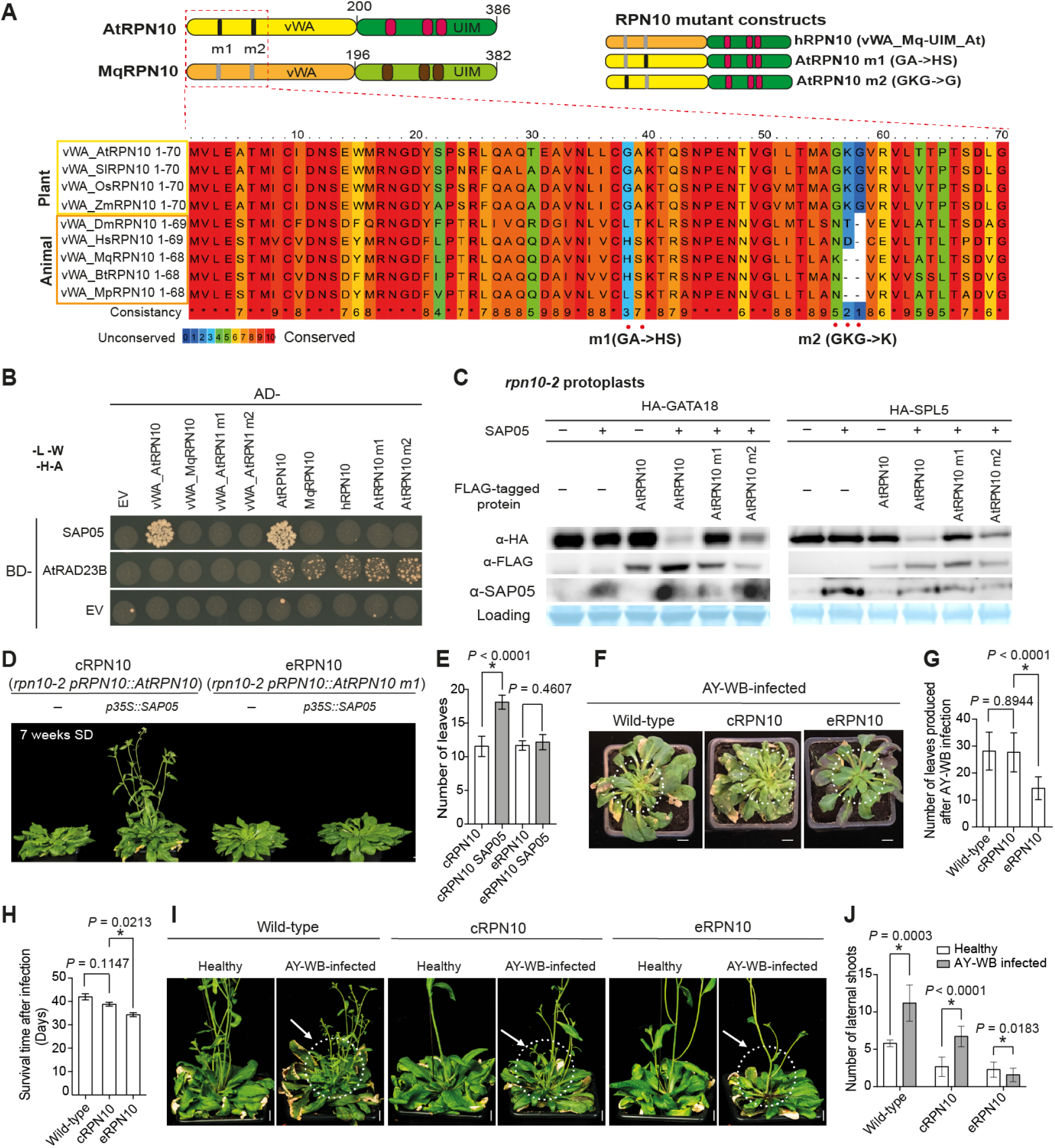
Insect-directed engineering of *A. thaliana* RPN10 confers resistance to SAP05 action. (A) Schematic overviews of domain organizations of *A. thaliana* and *M. quadrilineatus* RPN10 proteins and alignment of the first 70 residues of plant and animal RPN10 vWA domains. Highly divergent residues are highlighted below the alignment. Alignments of full-length RPN10 homologs are shown in Figure S6B. (B) Specific residues within the *A. thaliana* RPN10 vWA domain are required for SAP05 interaction in Y2H assays. See legends of Fig. 2 and Fig. 5A for abbreviations. Yeast growth on -L-W medium is shown in Fig. S10C. Interaction of the AtRAD23B proteasome shuttle factor with AtRPN10 is included as a control to show that RPN10 is functional as an interactor in yeast. (C) Specific residues within the *A. thaliana* RPN10 vWA domain are required for SAP05 degradation of plant GATA and SPL in *A. thaliana* protoplasts. See legend of Fig. 2E. GFP, control. (D-J) Specific residues within the *A. thaliana* RPN10 vWA domain are required for leaf and stem proliferations of *A. thaliana* plants in the presence of constitutively expressed SAP05 (D, E) and during AY-WB phytoplasma infection (F-J). *A. thaliana* plants included in these experiments were *rpn10-2* null mutants complemented with wild-type AtRPN10 (cRPN10) or AtRPN10 m1 (eRPN10). Scale bars, 1 cm. The symptomatic leaves in (F) and lateral shoots in (I) are circled. Phenotypes were statistically analyzed for number of leaves of 4-week-old plants (E), number of newly produced rosette leaves after AY-WB infection (G), plant survival time after AY-WB inoculation (H) and numbers of lateral shoots in control and infected plants (J). Data are mean ± s.d. from 2 independent experiments. *, *P* < 0.05, two-tailed unpaired Student’s *t-*test. See also Figure S8 for the leaf morphology of uninfected plants.

Stable transgenic *A. thaliana* plants that constitutively express genes of the SPL-interacting SAP05_WBLDa_ and SAP05_PnWBa_ phenotypically resembled the *miR156* overexpression plants or a high order *spl* mutant (*spl9 spl11 spl13 spl15*) (Figures 4E-H), including the production of more juvenile leaves (Figure 4G) and delayed flowering (Figure 4F), unlike the *MIM156* plants in which *miR156* activity is reduced via target mimicry (Franco-Zorrilla et al., 2007). In contrast, transgenic plants expressing GATA-interacting SAP05_WBLDb_ and SAP05_PnWBb_ did not show much difference in the production of juvenile leaves compared to wild type plants. Neither did a quadruple *gata* mutant, *gnc gnl gata17 gata17l*, nor the overexpression of a GATA member, *GNC*, that significantly impair plant greening and growth (Ranftl et al., 2016; Richter et al., 2010), altered the juvenile leaf number. However, phenotypes of SAP05_WBLDb_ and SAP05_PnWBb_ plants resembled those of well-characterised *gata* mutants in regarding to their early flowering (Figure 4E-F) (Ranftl et al., 2016; Richter et al., 2013; Richter et al., 2010), production of shorter siliques (Figure 4H) (Ranftl et al., 2016) and narrower rosette leaves with smooth margins (Figure S4) (Ding et al., 2015). Noticeably, SAP05 homologs that bind either SPLs or GATAs did not induce the full witch’s broom-like phenotypes that were observed in plants expressing the SAP05 homologs that bind both SPL and GATA transcription factors. This suggests that the concurrent destabilization of SPLs and GATAs by SAP05, decouples normal plant juvenile-to-adult and vegetative-to-reproductive developmental transitions, generating plants that retain juvenile characteristics and that nonetheless bolt to produce flowering shoots that remain sterile. Therefore, the combination of SAP05-mediated SPL and GATA degradation leads to the induction of the witch’s broom phenotype.

### Engineering plant RPN10 for resistance to SAP05 activity

Animals also encode GATA transcription factors and RPN10 proteins but lack SPLs. GATA transcription factors have multiple roles in humans and animals, including the regulation of processes related to immunity (Shapira et al., 2006). *SAP05* expression has been also detected in phytoplasma-carrier insects, in addition to infected plants (MacLean et al., 2011; Oshima et al., 2011). To investigate if phytoplasma SAP05 also interacts with GATA proteins of the leafhopper vector, we mined the transcriptome assembly of *M.quadrilineatus* (Drurey et al., 2019) for GATA transcription factors and identified six distinct transcripts with typical GATA ZnF domains (Figure S5A). However, none of these ZnF domains were shown to interact with SAP05 in Y2H assays (Figure S5B), indicating that SAP05 may not interact with insect vector GATA transcription factors.

We also identified a *M. quadrilineatus* RPN10 homolog (MqRPN10) that is highly similar in sequence to *A. thaliana* RPN10 (AtRPN10) (Figures 5A and S6). However, SAP05 of AY-WB did not interact with MqRPN10 in Y2H assays, nor with the MqRPN10 vWA domain, or a hybrid RPN10 (hRPN10) consisting of the *M. quadrilineatus* vWA domain and the *A. thaliana* C-terminal (UIM) domain (Figure 5B). Comparison of multiple vWA domains of plant and animal RPN10 homologues revealed differences between the two groups in two regions corresponding to amino acids 38–39 (GA vs. HS) and 56–58 (GKG vs. K--) in the *A. thaliana* vWA domain (Figure 5A). Altering these residues within *A. thaliana* RPN10 to those present in the *M. quadrilineatus* homologue, to create RPN10_38GA39->HS (m1) and RPN10_56GKG58->K (m2), resulted in a loss of SAP05 binding in Y2H assays (Figure 5B). The RPN10 variants interacted with the *A. thaliana* RADIATION SENSITIVE23 (RAD23B) protein, a ubiquitin shuttle factor that binds RPN10 UIM domains (Farmer et al., 2010), indicating that the RPN10 variants are functional in the Y2H assays. In addition, SAP05 degradation assays of AtGATA18 and AtSPL5 in *A. thaliana rpn10-2* protoplasts showed that these SAP05-targtes were less degraded in the presence of AtRPN10 m1 compared to AtRPN10 or AtRPN10 m2 (Figure 5C), indicating that the AtRPN10 vWA domain, and particularly the 38GA39 residues that are unique to plant versus animal RPN10 proteins, are involved in SAP05 binding and SAP05-mediated degradation of AtGATA18 and AtSPL5. In support of this notion, plant SUMO-tagged RPN10 vWA domains that carried the 38GA39->HS mutations (SUMO-evWA) had lower affinity for SAP05 in *E. coli* lysates compared to wild-type RPN10 vWA domain (Figure 3E), and the His-SUMO-tagged evWA domain did not pull down ZnF domain of AtSPL5 in the presence of SAP05, whereas His-SUMO-tagged vWA did (Figure 3F). Therefore, 38GA39 is involved in mediating the direct binding of SAP05 of AtRPN10 *in vitro and in vivo*.

In light of the above finding, we considered the possibility of engineering plant RPN10 as a way to block SAP05 activities. Noticeably, RPN10 has an essential function in *A. thaliana*, and *rpn10* null mutants have pleotropic phenotypes, including severe growth defects (Lin et al., 2011; Smalle et al., 2003). However, we found that introduction of the *A. thaliana* RPN10 mutant *rpn10-2* with the SAP05 non-interacting allele *AtRPN10 m1* under the control of the native RPN10 promoter to create *rpn10-2 pAtRPN10::AtRPN10 m1* plants (for simplicity, henceforth referred to as eRPN10 (engineered RPN10) plants), largely rescued the developmental defects of the *rpn10-2* plants. The eRPN10 plants did not have obvious phenotypic differences when compared to the *rpn10-2* complemented with a wild-type *A. thaliana* RPN10 allele (*rpn10-2 pAtRPN10::AtRPN10* plants; henceforth referred to as cRPN10 (complementation RPN10) plants) (Figure S7). Introducing the *p35S::SAP05* construct into cRPN10 plants yielded typical SAP05 phenotypes in 30 out of 34 independent transformants. In contrast, introduction of the *p35S::SAP05* construct in eRPN10 background generated wild type looking plants without obvious developmental phenotypes for all 33 independent transformants (Figures. 5D and 5E). Therefore, the RPN10_38GA39->HS mutation confers resistance to phytoplasma-SAP05-mediated developmental changes in *A. thaliana*.

To investigate the contribution of SAP05 to symptom development due to AY-WB phytoplasma infection in *A. thaliana*, we infected wild-type, cRPN10 and eRPN10 plants with AY-WB phytoplasma. The infected wild-type and cRPN10 plants produced increased quantities of small, deformed leaves and more lateral shoots compared to plants of similar age not infected with phytoplasma (Figures 5F-J and S8). The symptoms of these infected plants resembled the phenotypes of *p35S::SAP05* (Figures 1A-1C) and cRPN10 *p35S::SAP05* plants (Figure 5D). By contrast, eRPN10 *A. thaliana* infected with phytoplasma did not produce severely deformed leaves nor an increased number of lateral shoots as compared to the non-infected plants (Figures 5F-J). Moreover, the leaves of infected eRPN10 *A. thaliana* plants showed enhanced reddening compared to the wild-type and cRPN10 plants (Figures 5F and S8), indicating that SAP05 actions may reduce the plant stress-induced senescence during phytoplasma infection. Indeed, AY-WB infected eRPN10 plants died earlier compared to both cRPN10 and wild type plants (Figure 5H). All AY-WB-infected *A. thaliana* genotypes produced leaf-like flowers (Figure S9) that resemble the phyllody symptoms of AY-WB-infected plants, indicating that the engineered RPN10 allele does not interfere with the leaf-like flower phenotype induced by another AY-WB phytoplasma effector, SAP54 (MacLean et al., 2014; MacLean et al., 2011). These data demonstrate that the AY-WB SAP05 effector is largely responsible for the shoot proliferation /witch’s-broom-like symptoms during AY-WB infection of *A. thaliana* and the blocking of SAP05 activities reduces the host tolerance towards this phloem-inhabiting insect-vectored bacterial pathogen.

## Discussion

This work shows that SAP05 effector co-opts the plant 26S proteasome ubiquitin receptor RPN10 to mediate degradation of SPL and GATA—two distinct classes of plant transcription factors—through a lysine ubiquitination-independent process (Fig. 6A). SPL transcription factors have a conserved role in controlling developmental phase transitions of vascular plants (Huijser and Schmid, 2011; Wang et al., 2009; Xu et al., 2016), whereas GATA transcription factors regulate plant organ development, timing of flowering and branching patterns in both dicots and monocots (Ding et al., 2015; Hudson et al., 2013; Ranftl et al., 2016; Richter et al., 2010; Zhang et al., 2013). Phenotypes induced by phytoplasma SAP05 are consistent with the simultaneous degradation of SPLs and GATAs as this would lead to decoupling of plant developmental phase transitions and subsequent delayed plant aging and promotion of witch’s broom-like excessive vegetative tissue and sterile adult shoot production (Figure 6B). Our model describes a mechanistic framework for how obligate parasites can induce complex and substantial phenotypic changes in their hosts in ways that favour their transmission to other trophic levels.

**Figure 6.**
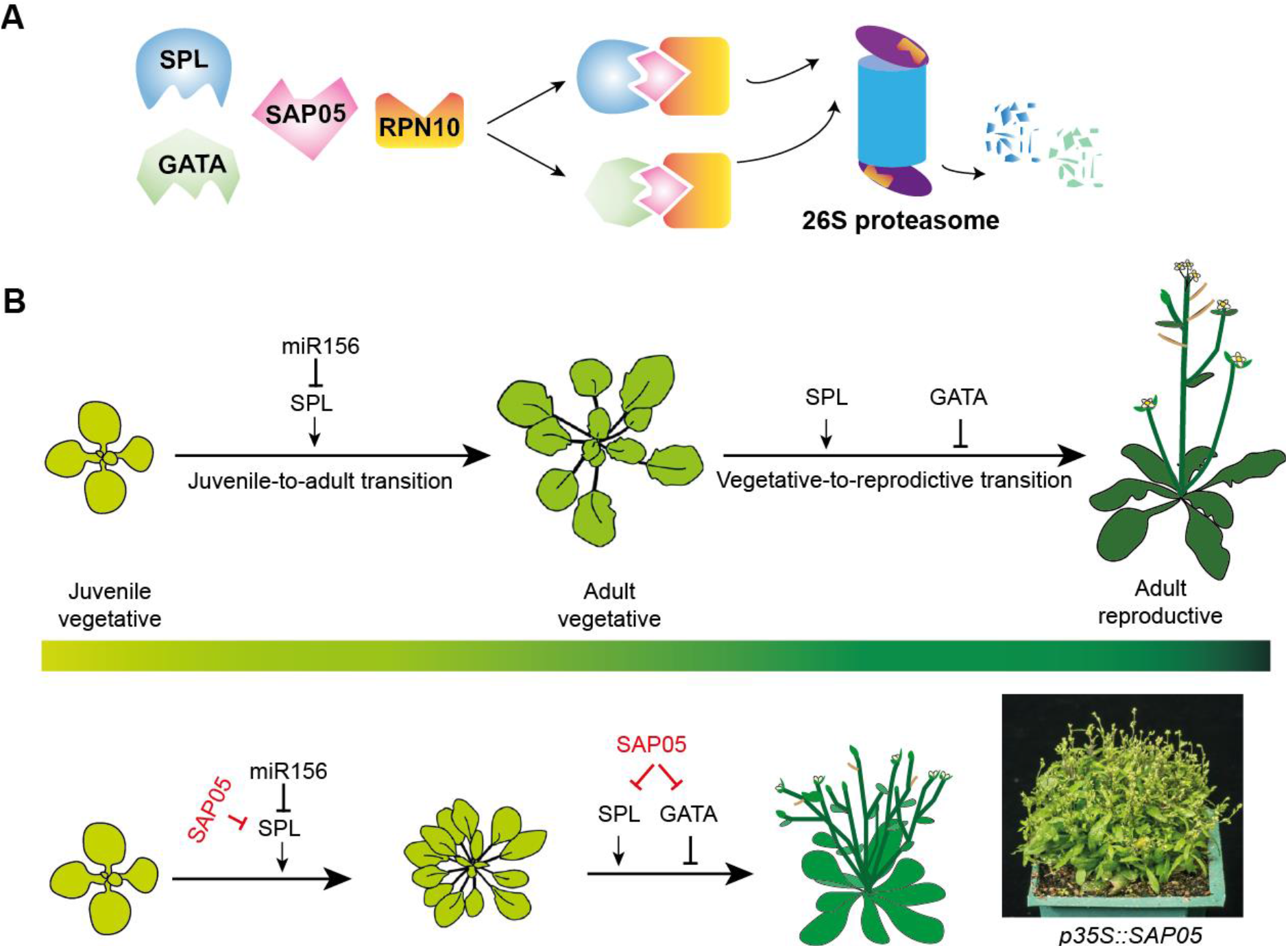
Model for SAP05-induced delayed aging and witch’s broom formation. (A) A schematic overview of SAP05-mediated selective protein degreadation in the plant host 26S proteasome. (B) During normal plant growth, *SPL* genes are regulated by *miRNA156* to ensure the proper progression of plant developmental phase transitions while GATA genes regulate multiple processes of plant development, including the suppression of flowering and branching (upper panel). In phytoplasma infected plants, phytoplasma effector SAP05 mediates, on the one hand, the destabilization of SPL transcription factors generating juvenized plants, and on the other hand, the destabilization of GATA proteins thereby releasing their flowering and branching inhibition. As a whole, the uncoupling of plant developmental phase transitions by phytoplasma SAP05 contributes to witch’s broom formation.

Phytoplasmas are restricted to the phloem. These bacteria benefit from the SAP05-induced phenotypes by increasing numbers of leaves and adult shoots to generate more vascular tissues that phytoplasmas can colonise. In support of this notion, the engineered RPN10 plants which are resistant to SAP05 activities, do not produce more vegetative tissues and die earlier during phytoplasma infection (Figures 5F-J). Moreover, whereas flower structures are formed, these remain sterile. So photosynthates transported within the phloem are not being used for flowering and seed production, and may instead be consumed by the phytoplasmas, which are known to accumulate in plant sink tissues. Enrichments of nutrients are likely important, because phytoplasmas lack essential metabolic pathways and depend on the import of a diverse range of metabolites, including sugars, nucleotides, amino acids and ions, mostly via ABC transporters (Bai et al., 2006; Oshima et al., 2004). Finally, an obvious advantage for the obligate phytoplasmas to delay plant senescence and death is that it can increase the chance of disease transmission by the insect vectors that also feed on the plant hosts. Therefore, the SAP05-induced decoupling of host developmental transitions likely promotes plant developmental traits that facilitate survival and spread of this ‘Zombie plant’ pathogen.

Our studies led to a strategy for engineering plants to be insensitive to SAP05 activities. Despite RPN10 being an extremely conserved protein among eukaryotes, including plants and arthropods, SAP05 does not bind RPN10 of the leafhopper vector. The SAP05-binding specificity to RPN10 is dependent on just two amino acids that, fascinatingly, are one of the only few lowly conserved sequences between plant and animal RPN10 vWA domains (Fig. 5A). Introduction of these insect vector vWA amino acids into the plant vWA domain generated a functional plant RPN10 and plants that are more resistant to SAP05 activity during phytoplasma infection. Phytoplasmas are invasive colonisers of their insect vector, as these bacteria enter gut cells of these insects and replicate in their gut epithelia, haemolymph and salivary glands (Koinuma et al., 2020). The findings that SAP05 does not bind leafhopper RPN10 homologs, nor insect GATA transcription factors, suggest that SAP05 effectors specifically interact with plant RPN10 and GATA proteins and may have evolved to avoid binding these proteins in insect vectors. Whether SAP05 effectors modulate processes in insect vectors remains to be investigated. Nonetheless, these data indicate that plant RPN10 is a plant susceptibility factor and Achilles’ heel for phytoplasmas. Hence, the introduction of single nucleotide changes in RPN10 genes, by for example CRISPR-Cas technologies (Chen et al., 2019), is a promising strategy to achieve a level of durable resistance of crops to phytoplasmas.

We found that SAP05 homologs of divergent phytoplasmas differentially interact with plant SPLs and GATAs. Whereas the majority of SAP05 homologs bind members of both the plant transcription factor families, some SAP05 homologs only bind and degrade either SPLs or GATAs. In the SAP05 phylogenetic tree, SAP05 homologs targeting both transcription factors are on short branches (Figure 4D) indicating low mutation rates of these effector genes, possibly because of constraints on mutations that are permitted to allow SAP05 interactions with both SPLs and GATAs. In contrast, the SAP05 that have evolved to only bind one of the transcription factor families have longer branches in the phylogeny (Figure 4D), indicating higher mutations rates of these *SAP05* genes that is consistent with directional selection leading to specialization that often occurs upon gene duplications within families (Taylor and Raes, 2004). Taken together, these data suggest that SAP05 is under adaptive selection to interact with both RPN10 and transcription factors of plants.

The way by which SAP05 bridges between transcription factors and a conserved proteasome component is unique. A plethora of parasites exploit the host cell ubiquitin machinery to promote degradation of target proteins and subvert host processes (Ashida et al., 2014; Banfield, 2015; Lin and Machner, 2017). Effectors described so far interfere with aspects of the ubiquitination machinery, such as ubiquitin (Cui et al., 2010) and E1, E2 and E3 ligase activities (Abramovitch et al., 2006; Qiu et al., 2016; Sanada et al., 2012; Tzfira et al., 2004), or inhibit the general enzymatic activities of the host 26S proteasome (Groll et al., 2008; Ustun et al., 2013). However, to date, effectors such as SAP05 that directly link host targets to the proteasome system to mediate degradation of targets in a manner that does not require lysine ubiquitination of the substrates are not known. This SAP05 mode of action is potentially significant. Whereas cellular protein levels may be altered by gene knockout and RNA silencing, some systems require the direct targeting of proteins for degradation. In fact, targeted protein degradation has become one of the most promising approaches for drug discovery in targeted therapies (Chamberlain and Hamann, 2019). Current approaches to changing protein abundance in cells rely on substrate ubiquitination: for example, the proteolysis-targeting chimera (PROTAC) technique uses small-molecule ligands that create complexes between E3 ligases and targets, a process that can be challenging (Schapira et al., 2019). Phytoplasma SAP05 effectors may enable a more direct way to mediate degradation of target proteins and given that SAP05 effectors differ in binding affinities to SPLs and/or GATAs may be engineered to degrade other proteins.

In contrast to most pathogens effectors that target immune responses of their hosts, this and previous works show that phytoplasmas effectors have converged onto modulating key plant developmental regulators. These include plant TCP transcription factors (Sugio et al., 2011a), the MADS-box transcription factors (MacLean et al., 2014; Maejima et al., 2014), and here, the SPL and GATA transcription factors. Members of these transcription factors regulate each-other at overlapping and different plant developmental stages. Phytoplasma effector perturbation of large protein families rather than individual members is likely beneficial for exerting control over multiple host processes simultaneously, a strategy similar perhaps to the targeting of protein families by microRNAs (Samad et al., 2017). Moreover, because phytoplasmas are being introduced into plants at different stages of plant development, depending on when their insect vectors feed, it is likely beneficial for the obligate parasites to be able to control plant developmental processes throughout the plant age stages. The latter are characterized by juvenile-to-vegetative and vegetative-to-adult developmental stage transitions. The finding that SAP05 effectors mediate degradation of SPLs and GATAs that control precisely these transitions (Fig. 6B) is therefore particularly likely to enable phytoplasma to take control of their plant host and corroborates the view that the developmental changes enforced by the multi-host obligate phytoplasma parasites are adaptive.

## Materials and Methods

### Plant materials and growth conditions

*A. thaliana* Columbia-0 ecotype (*Col-0*) plants were grown in the greenhouse under either long-day (16 h light/8 h dark) or short-day conditions (10 h light/8 h dark) at 22°C. Transgenic plants were generated as previously described (MacLean et al., 2011). For generating *p35S::SAP05* or *pAtSUC2::SAP05* plants, codon-optimised SAP05 coding sequences (without the secretory signal peptide) were used. SAP05 sequences used for generating transgenic plants were listed in Table S4. Plasmids used in this study are listed in Table S5.

### Plant phenotypic analysis

Plant age was determined from the date seeds were transferred to growth chambers. Juvenile leaves refer to rosette leaves that only produce trichomes on their adaxial side. Rosette leaves that have trichomes on both side of leaves (adaxial and abaxial) were recorded as adult leaves. Bolting time was recorded when the main inflorescence reached a height of 0.5 cm.

### Yeast two-hybrid analysis

The initial Y2H screen of SAP05 against the *A. thaliana* seedling library was performed by Hybrigenics Services SAS (Paris, France). The coding sequence of SAP05 without the secretory signal peptide was cloned into a pB27 bait plasmid as a C-terminal fusion to the LexA domain (Table S1). The prey library was constructed from an *A. thaliana* seedling cDNA library, with pP6 as the prey plasmid. In a second yeast two-hybrid screen, the same SAP05 sequence was cloned into the pDEST32 plasmid and screened against an *A. thaliana* transcription factor library (pDEST22-TF) (Pruneda-Paz et al., 2014). The identified interactions were further confirmed using the Matchmaker Gold yeast two-hybrid system (Clontech) or the DUALhybrid system (Dualsystems Biotech). SPL and GATA proteins identified in these screens are summarised in Table S2.

### Protoplast degradation assays

*A. thaliana* (Col-0) mesophyll protoplast isolation and transformation were carried our as reported (Yoo et al., 2007). Briefly, mesophyll protoplasts were isolated from leaves of 4–5-week-old *A. thaliana* plants grown under short-day conditions. For transfection, 300 μl of fresh protoplast solution (120,000 protoplasts) was transformed with 24 μg of high-quality plasmids (12 μg each for co-transfection) using the PEG-calcium method. Transfected protoplasts were incubated at room temperature (22-25 °C) for 16 h in the dark before harvest. For drug treatment, a final concentration of 20 μM MG132 (Sigma), 5 μM Bortezomib (Sigma) or 10 μM E-64d (Sigma) or 5 mM 3-MA (Sigma) were added during the 16-h incubation period. Except for 3-MA which was prepared as 0.1 M stock solution in water, the others were prepared as 10 mM stock solution in DMSO. Equivalent volume of DMSO was used as mock control. For detection of proteins on western blots, whole protein extracts from protoplasts were separated on NuPAGE 4-12% Bis-Tris Gels (Invitrogen) and transferred to 0.45-μm PVDF membranes (Thermo Scientific) using the Bio-Rad mini-PROTEAN Electrophoresis system. Membranes were blocked by incubation in 5% (w/v) milk power in phosphate-buffered saline and 0.1% (v/v) Tween-20 for 2 h at room temperature. Primary antibody incubation was carried out at 4 °C overnight. Antibody to SAP05 from AY-WB phytoplasma were raised to the mature part of the SAP05 protein (residues 33–135), which was produced with a 6XHis-tag into *Escherichia coli* and purified. The purified protein was used for raising polyclonal antibodies in rabbits (Genscript). Optimal detection of SAP05 in phytoplasma-infected plants occurred at a 1:2,000 dilution of the antibody, and this dilution was used in all western blot experiments for detection of SAP05. The OptimAb HA.11 monoclonal antibody (Eurogentec) was used to detect hemagglutinin (HA)-fusion proteins at the concentration of 0.5 μg/ml. The ANTI-FLAG monoclonal antibody (Sigma, F-3165) was used to detect FLAG tag-fusion proteins at a 1: 5000 dilution. Rabbit polyclonal anti-GFP antibody (Santa Cruz Biotechnology) was used at a 1:10,000 dilution. Protein loading was visualized using Amido black staining solution (Sigma).

### GUS staining and real-time PCR

The expression of *rSPL11-GUS* and *rSPL13-GUS* reporter genes and the GUS staining of healthy or phytoplasma-infected plants at 4 weeks after phytoplasma inoculation was performed as described previously (Xu et al., 2016).

### Protein expression, purification and in vitro binding assays

DNA that code for SAP05 (Ala33-Lys135), ZnF_AtSPL5 (Ser60-Leu127), vWA domian of AtRPN10 (Val2-Gly193) or a vWA mutant (38GA39->HS) were subcloned to either pOPINF (for N-terminal His tag), pOPINA (no tag) or pOPINS3C (for N-terminal His-SUMO tag) (Berrow et al., 2007). The vectors were transformed or co-transformed into *E.coli* strain BL21 (DE3). Protein expression and affinity purification using immobilized metal affinity chromatography (IMAC) were carried out according to manufacture’s instruction (Ni-NTA agarose, Qiagen). Briefly, protein expression was induced by the addition of 1 mM Isopropyl-β-D-thiogalactoside (IPTG) at 16°C for 20 h with shaking at 220 rev min^−1^. Cell pellets were lysed in IMAC buffer (50 mM Tris-HCl, 50 mM glycine, 0.5 M NaCl, 20 mM imidazole, 5% glycerol, pH 8.0) for affinity purification and eluted with elution buffer (50 mM Tris-HCl, 50 mM glycine, 0.5 M NaCl, 0.5 M imidazole, 5% glycerol, pH 8.0). Further purification was achieved by gel filtration (AKTA avant chromatography system) in gel filtration buffer (20 mM HEPES, 0.15 M NaCl, pH 7.5). When nessassay, the tags were removed by HRV 3C protease in gel filtration buffer. For testing complex formation in gel filtration, equal amount (molecular weight) of proteins were mixed in gel filtration buffer and left on ice for 45 mins before sample injection. For *in vitro* pull-down assay, His-tagged vWA or evWA domains were bound to 50 μL Ni-NTA agarose beads during protein purification starting from 50 mL cell culture and the beads were washed sequencially with IMAC buffer and gel filtration buffer. 20 μM SAP05 or/and ZnF domain of AtSPL5 were incubated with the Ni-NTA agarose beads in 100 μL gel filtration buffer for 1 h. After discarding the supernantant, the beads were wahed sequencially with gel filtration buffer and IMAC buffer. Proteins bound to the beads were eluted in 100 μL elution buffer. Input samples and pull-down samples were analysed by SDS-PAGE and coomassie staining. The experiments were repeat for 3 times with consistent results.

### Phytoplasma infection assays

*M. quadrilineatus* colonies carrying the AY-WB phytoplasma were reared on infected lettuce and china aster under long-day conditions at 24 °C. For *A. thaliana* inoculation, one leaf from a 4-week-old plant grown under short-day conditions was exposed to two or three leafhoppers from this colony in a clip cage for 2 d. The leaf and the clip cage were then removed to get rid of the carrier insects. For disease symptom recording, healthy or phytoplasma-exposed plants were then kept in short-day for observing leaf development and survival or transferred to long-day conditions for examining branching phenotypes.

### Phylogenetic analysis

Sequences were aligned with MUSCLE (v3.8.31) configured for highest accuracy. Phylogenetic trees were reconstructed using the maximum-likelihood method implemented in the PhyML program (v3.1/3,0 aLRT). Graphical representation and editing of the phylogenetic trees were performed with TreeDyn (v198.3).

### Statistical Analysis

Statistical analysis was performed in Prism 7. One-way ANOVA was used to analyse experimental data with more than 2 two experimental groups followed by Tukey’s multiple comparisons test, and two-tailed unpaired Student’s *t*-test was used for other data analysis.

## Acknowledgements

We thank Xiaolan Zhang (China Agricultural University) for the *han-2* and *han-2 p35S::CsHAN1* seeds, Claus Schwechheimer (Technical University of Munich) for *gnc gnl gata17 gata17l* and *p35S::GNC* seeds. We are grateful to the John Innes Centre (JIC) Horticultural Services for growing the plants, to the JIC Entomology Facility for rearing of leafhopper and phytoplasma stocks and to JIC Photographic Services for imaging experimental materials. We gratefully acknowledge summer student Luke Sherwin for technical support. The project was supported by the Human Frontier Science Program RGP0024/2105 (S.A.H. and R.G.H.I.) and a Marie Curie International Incoming Fellowship FP7-PEOPLE-2010-IIF Grant Agreement Number 274444 (A.M.M.). Additional support was received from Biotechnology and Biological Sciences Research Council grants BB/K002848/1, BB/J0045531/1 and BB/P012574/1 and the John Innes Foundation.

## Author contributions

S.A.H. conceived the project and S.A.H., R.G.H.I. and A.M.M. obtained funding. W.H. and S.A.H. designed the approach and experiments and W.H. conducted the majority of the experiments and data analyses described in the paper. A.S. and A.M.M. cloned SAP05 and performed initial phenotypic analysis of SAP05 plants. A.M.M. initiated the initial yeast two-hybrid experiments to identify SAP05 targets and purified SAP05 to raise antibodies to this protein. R.G.H.I. and M.B. performed yeast two-hybrid experiments to assess SAP05–plant transcription factor interactions. A.M. and S.K. provided input for the gel infiltration experiment to study *in vitro* protein-protein interactions. C.K. and S.C. contributed to the phylogenetic analysis of SAP05 homologs. W.H. and S.A.H. wrote the manuscript and all authors edited the manuscript.

## Data availability

All data generated or analyzed during this study are included in this published article and its Supplementary Information files. All materials within the paper are available from the corresponding author upon reasonable request.

---

**Figure S1.**
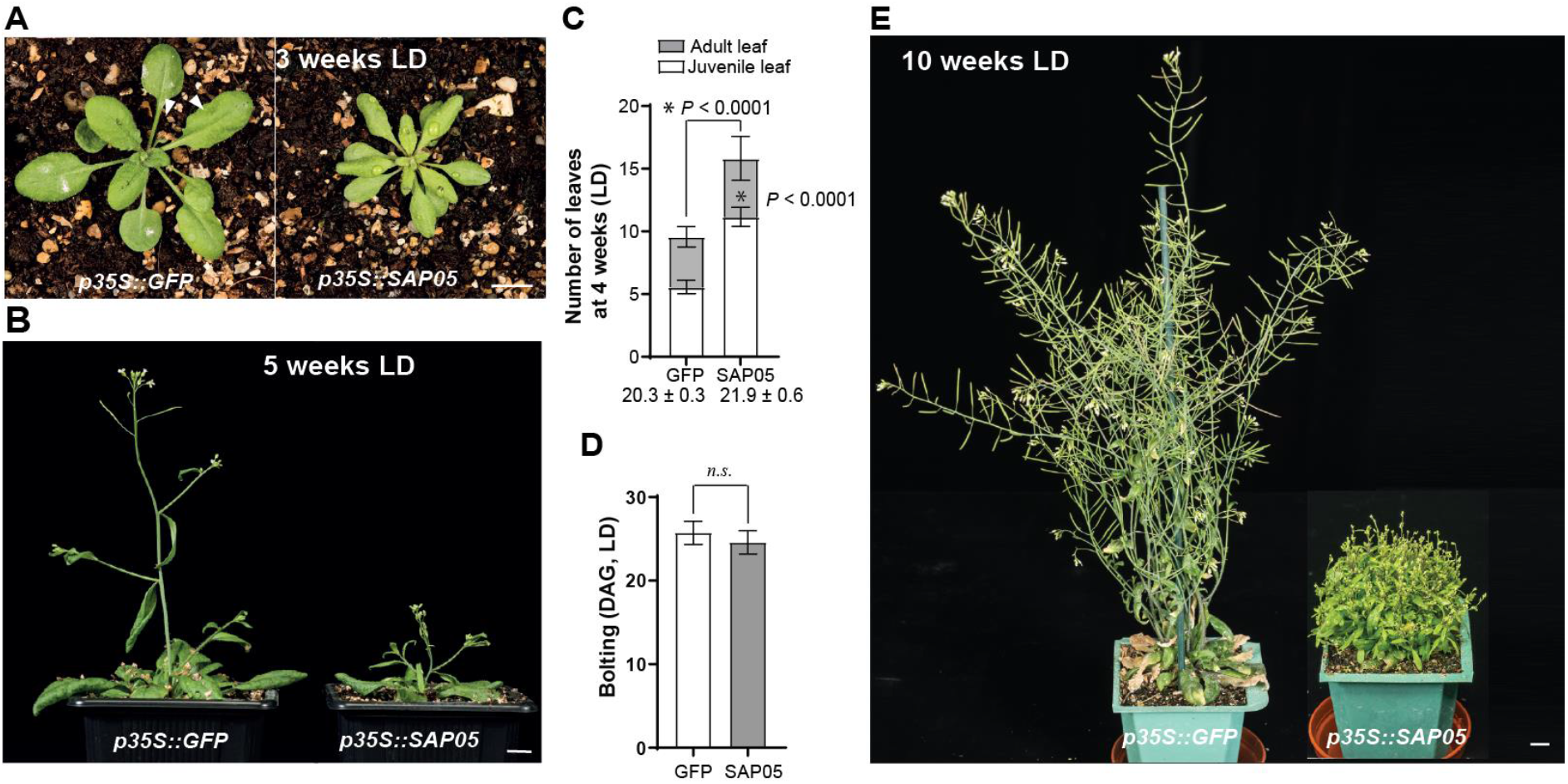
The morphology of SAP05-expressing *A. thaliana* under long-day conditions. (A-B) Representative images of plants stably producing SAP05 or GFP (control) grown under long-day (LD) conditions. Images were obtained at 3 weeks (A) and 5 weeks (B) after germination. Arrowheads in (A) indicate leaf serrations of GFP plants as opposed to the smoother leaf edges of SAP05 plants. Scale bars, 1 cm. (C-D) Statistical analysis of phenotypes shown in (A-B): numbers of rosette leave of 4-week-old plants (C) and time of shoot emergence from rosettes (bolting time; D). Numbers under the bars in (C) indicate the time (DAG) when the first abaxial trichome appeared. DAG, day after germination. *, *P* < 0.05, two-tailed unpaired Student’s *t*-tests. (E) A SAP05 plant exhibiting severe bushy and sterile phenotypes.

**Figure S2.**
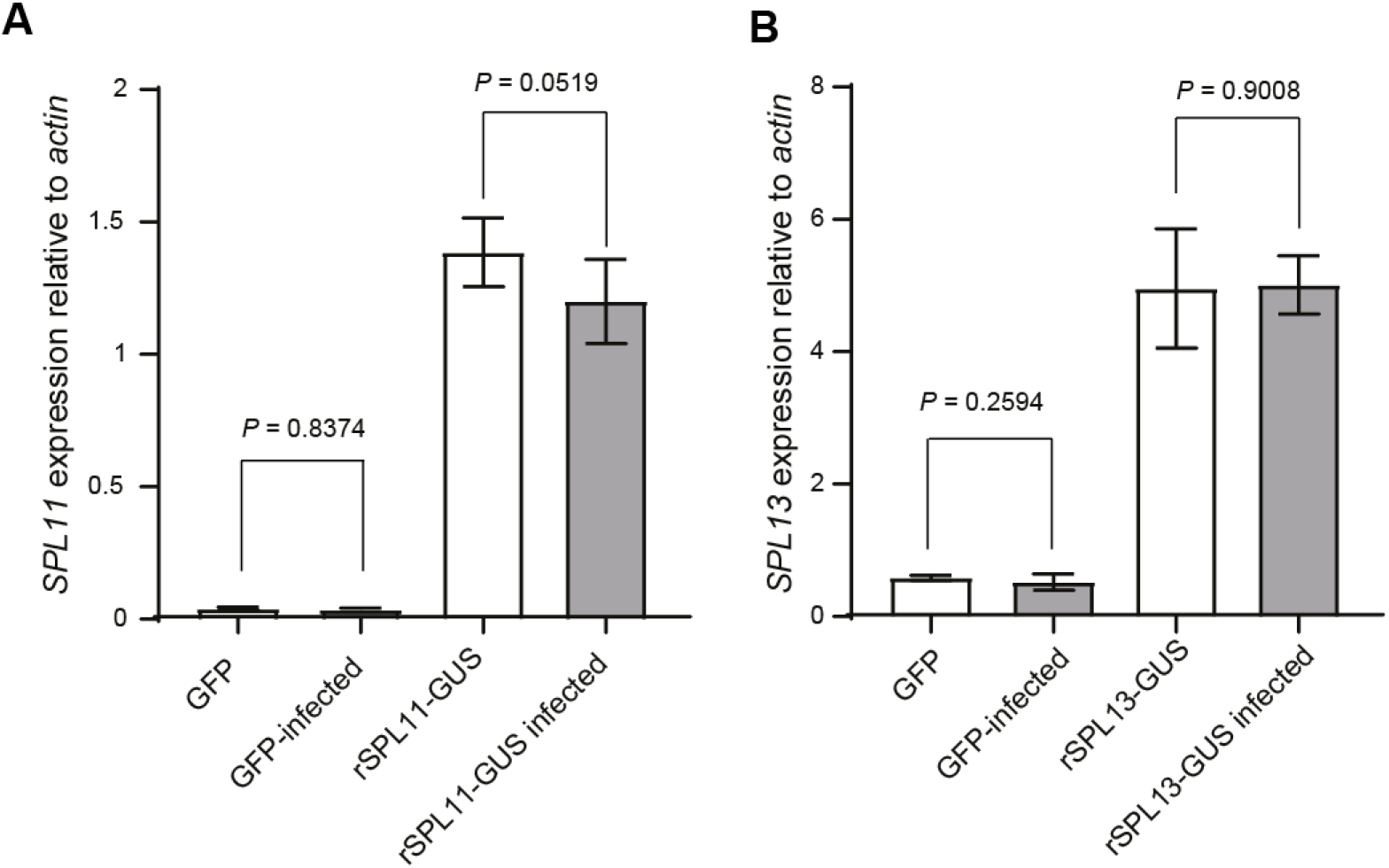
AY-WB phytoplasma infection does not change SPL genes expression. (A and B) The relative expression levels of *AtSPL11* (A) and *AtSPL13* (B) genes in healthy and AY-WB phytoplasma-infected control (GFP) or overexpression plants (rSPL11-GUS and rSPL13-GUS). The expression levels were shown as fold changes relative to the β actin gene. No significant difference was observed in each genotype, two-tailed unpaired Student’s *t-*tests.

**Figure S3.**
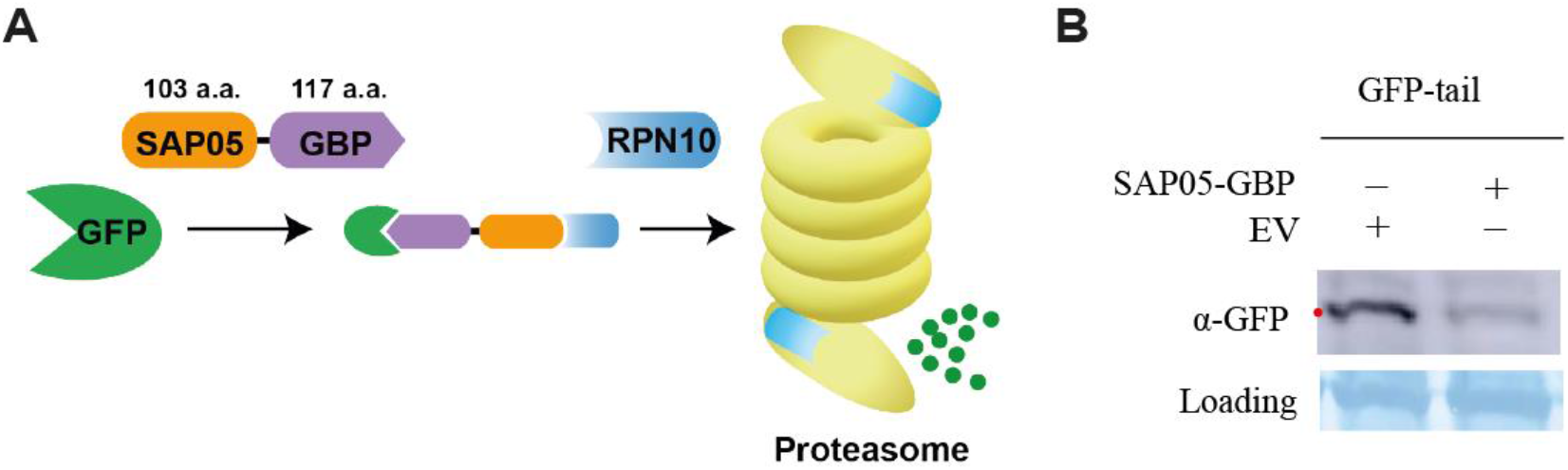
A SAP05-GBP fusion protein mediates the destabilization of GFP in A. *thaliana* protoplast. A. Proposed model for SAP05-mediated GFP degradation. The GBP (GFP-binding protein) derived from a single-chain antibody domain specifically recognising GFP was fused to SAP05 at its C-ter via a Glycine-rich linker. B. Western blot analysis of GFP abundance in the presence of SAP05-GBP or an empty vector control when transiently expressed in *A. thaliana* protoplasts. ‘tail’ represents an unstructured region that serves as an initiation site for proteasomal degradation.

**Figure S4.**
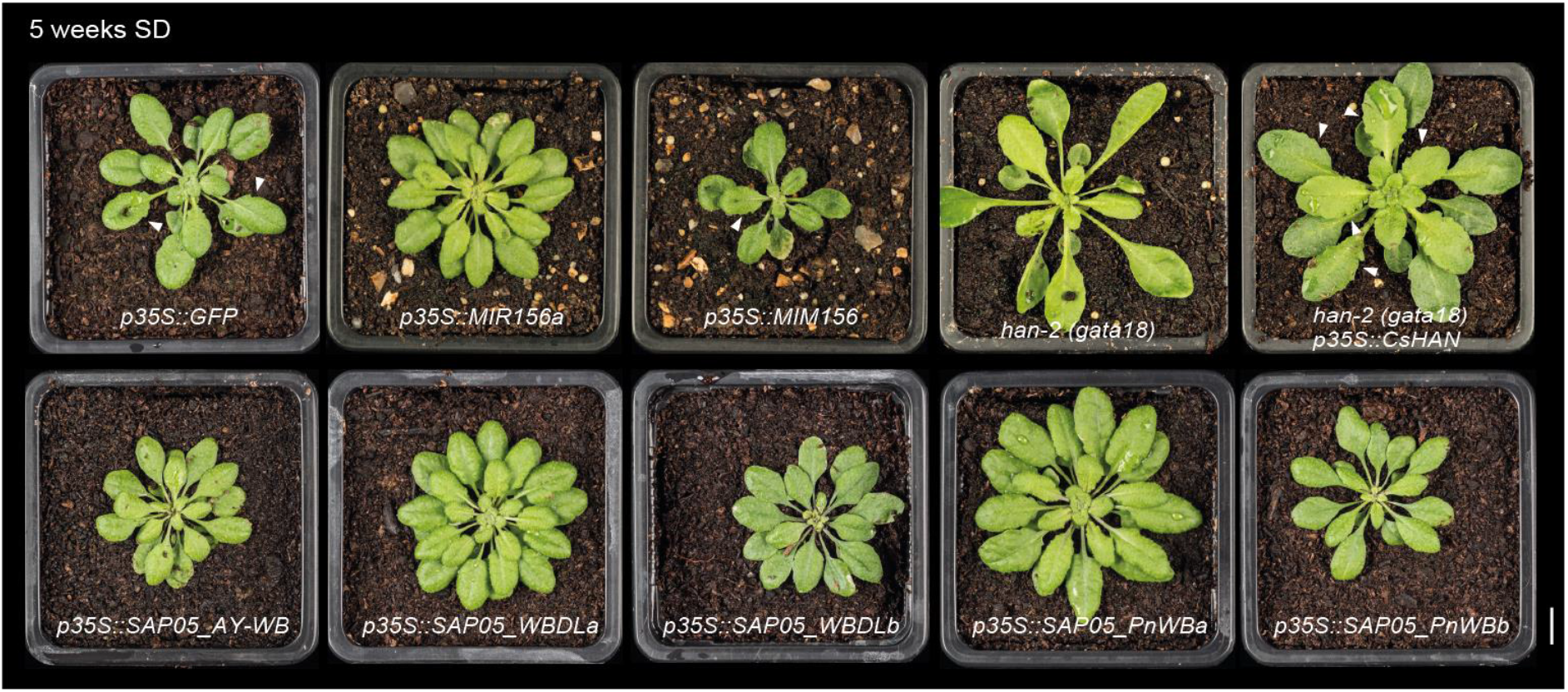
Different SAP05 homologs induce distinct leaf morphologies in *A. thaliana* reminiscent of either *MIR156* overexpression or a *GATA* mutant. Arrowheads indicate leaf serrations. The *A. thaliana han* mutant (*han-2*) produces rosettes with a smooth margin while the overexpression of a *Cucumis sativus L. GATA18* homologue (*CsHAN*) under the control of the 35S promoter in the *A. thaliana han-2* background leads to leaves with more severe serrations. Scale bar, 1 cm.

**Figure S5.**
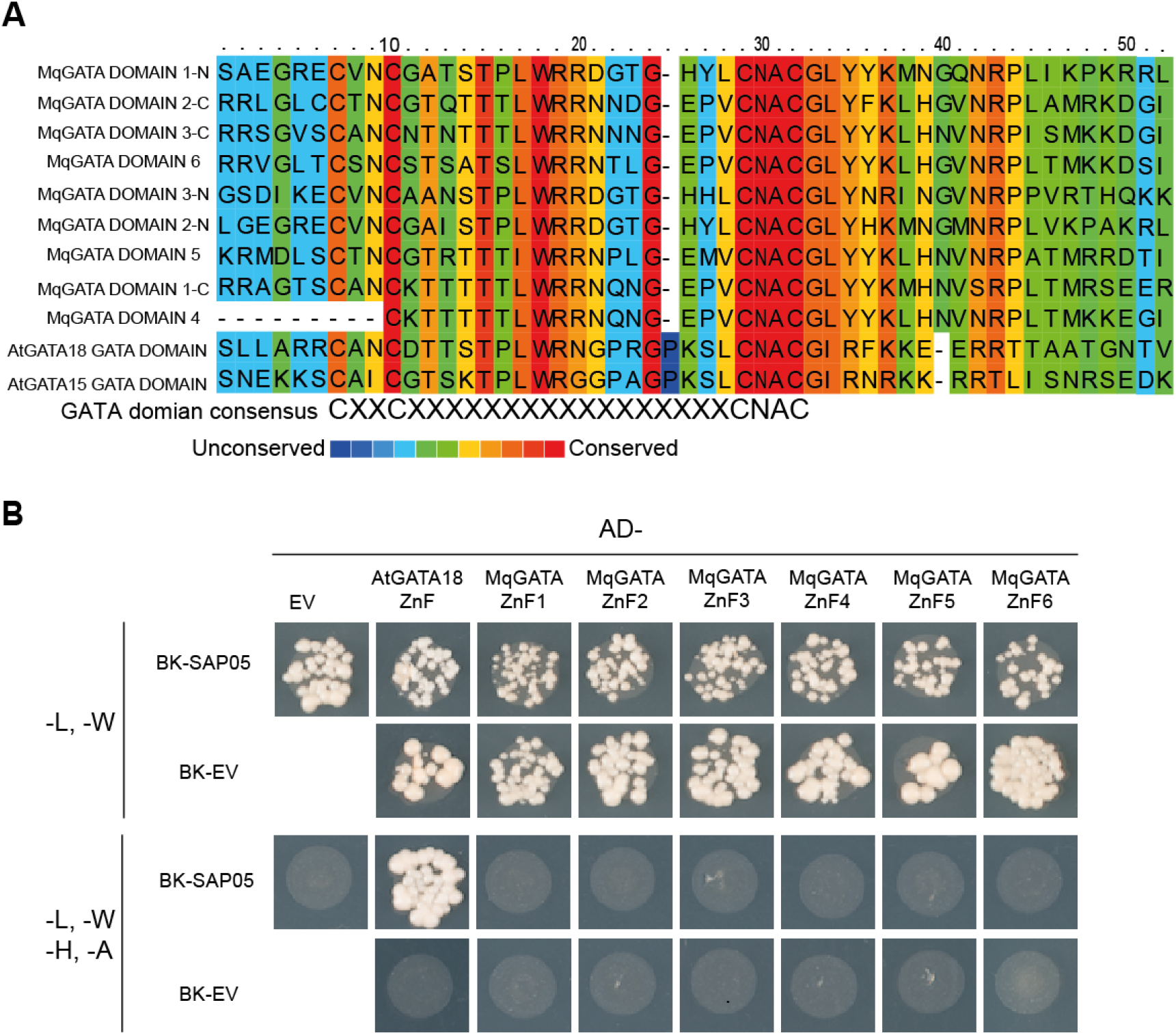
SAP05 does not interact with phytoplasma insect vector GATA domains in yeast two-hybrid assays. (A) Alignment of *M. quadrilineatus* GATA domains with two *A. thaliana* GATA domains. (B) Y2H analysis of SAP05 interaction with various *M. quadrilineatus* GATA domains. The GATA domain of AtGATA18 was used as a positive control.

**Figure S6.**
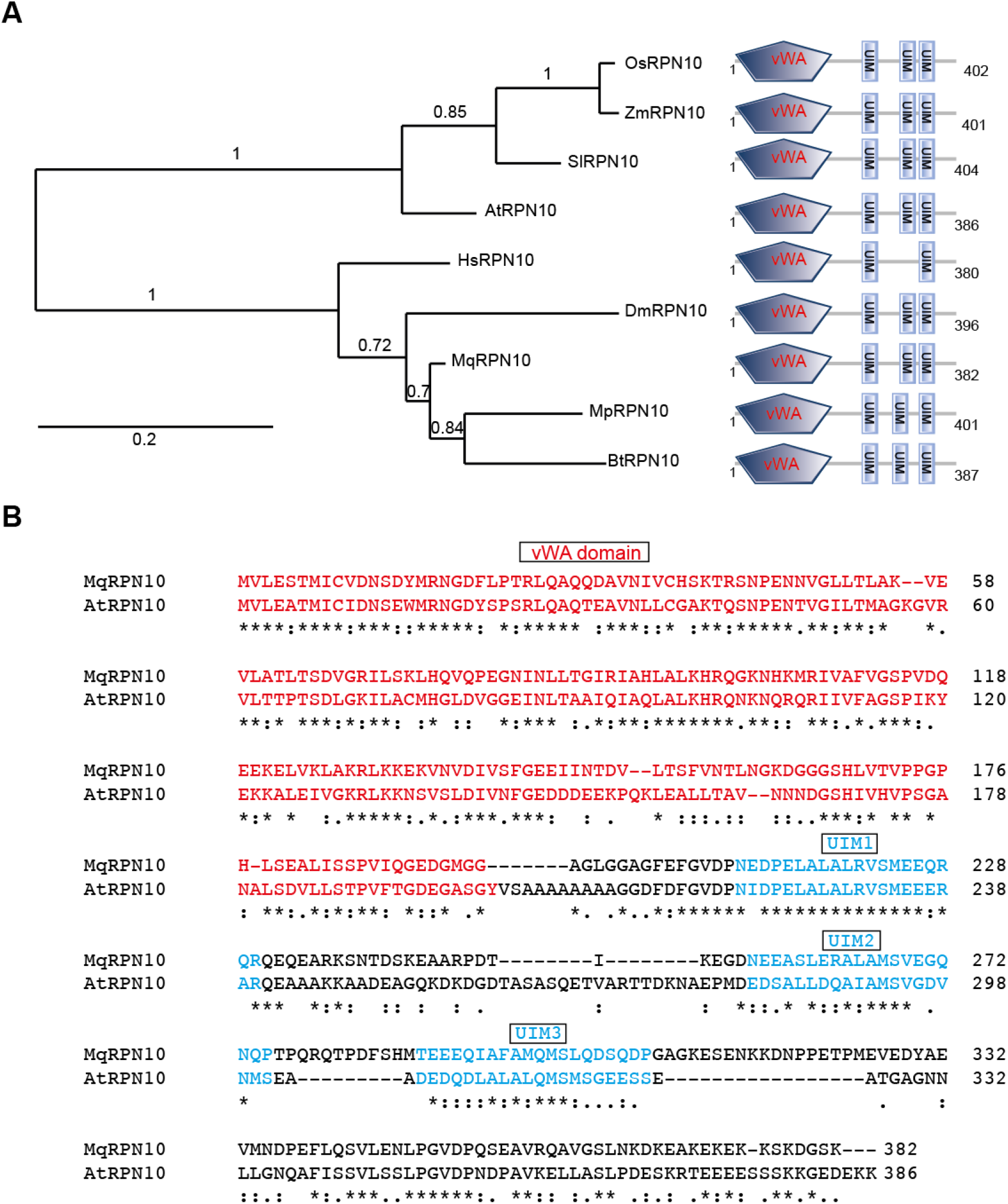
RPN10 homologues are conserved among plants and animals. (A) Phylogenetic analysis of RPN10 proteins from various organisms. The presence of vWA and UIM domains were predicted by PFAM. AtRPN10, *Arabidopsis thaliana* RPN10 (Uniprot ID: P55034); SlRPN10, *Solanum lycopersicum* RPN10 (A0A3Q7F6N7); OsRPN10, *Oryza sativa* RPN10 (O82143); ZmRPN10, *Zea mays* RPN10 (B6TK61); DmRPN10, *Drosophila melanogaster* RPN10 (P55035); HsRPN10, *Homo sapiens* RPN10 (Q5VWC4); BtRPN10, *Bemisia tabaci* RPN10 (XP_018915695); MqRPN10, *Macrosteles quadrilineatus* RPN10; MpRPN10, *Myzus persicae* RPN10 (XP_022181722.1). (B) Sequence alignment of the *A. thaliana* RPN10 and the *M. quadrilineatus* RPN10 proteins. The vWA domains and UIM domains are highlighted in red and blue, respectively.

**Figure S7.**
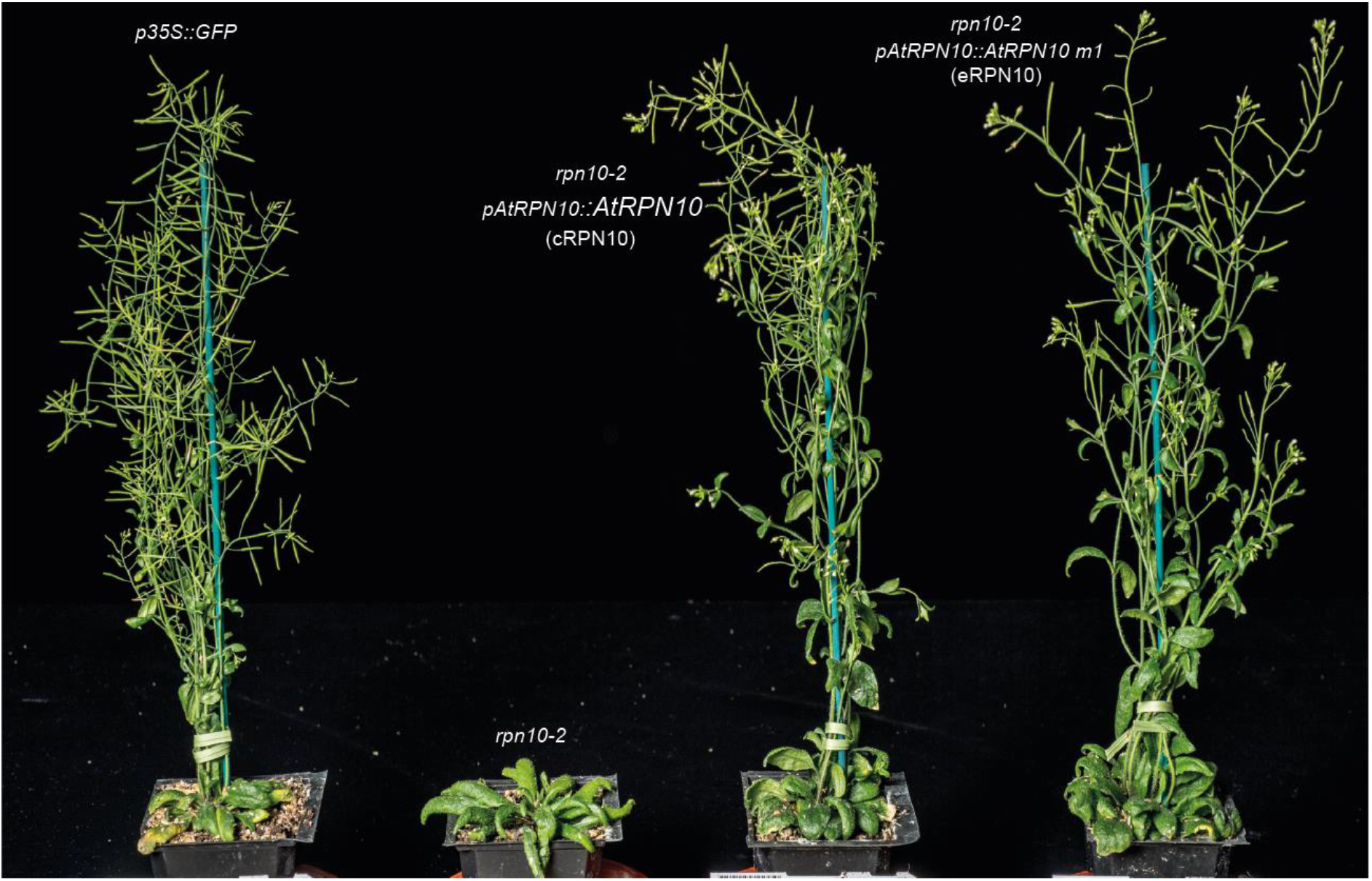
An engineered *RPN10* allele rescues the developmental defects of the *rpn10-2* mutant. The *rpn10-2* mutant was complemented by either a wild-type *AtRPN10* allele (cRPN10) or an engineered RPN10 allele (AtRPN10 m1, eRPN10) under the control of the native promoter. At least two independent lines for each complementation were obtained, with consistent plant phenotype.

**Figure S8.**
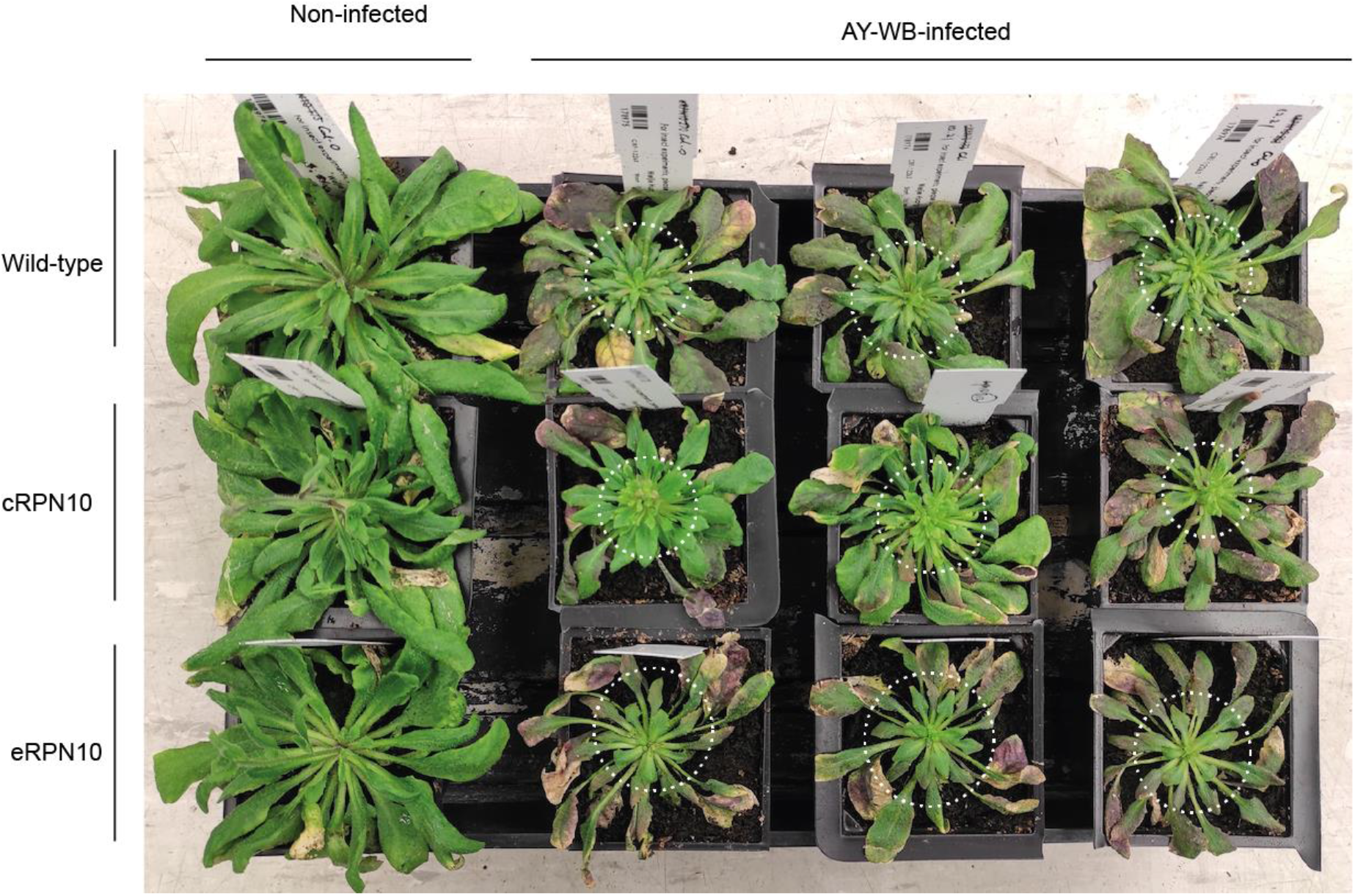
Rosette leaf morphology on healthy and AY-WB phytoplasma-infected plants. Representative images of healthy *A. thaliana* plants of different backgrounds and plants infected with AY-WB phytoplasma. All plants were kept in short-day conditions throughout the experiment. Circled areas correspond to rosette leaves that emerged during phytoplasma infection.

**Figure S9.**
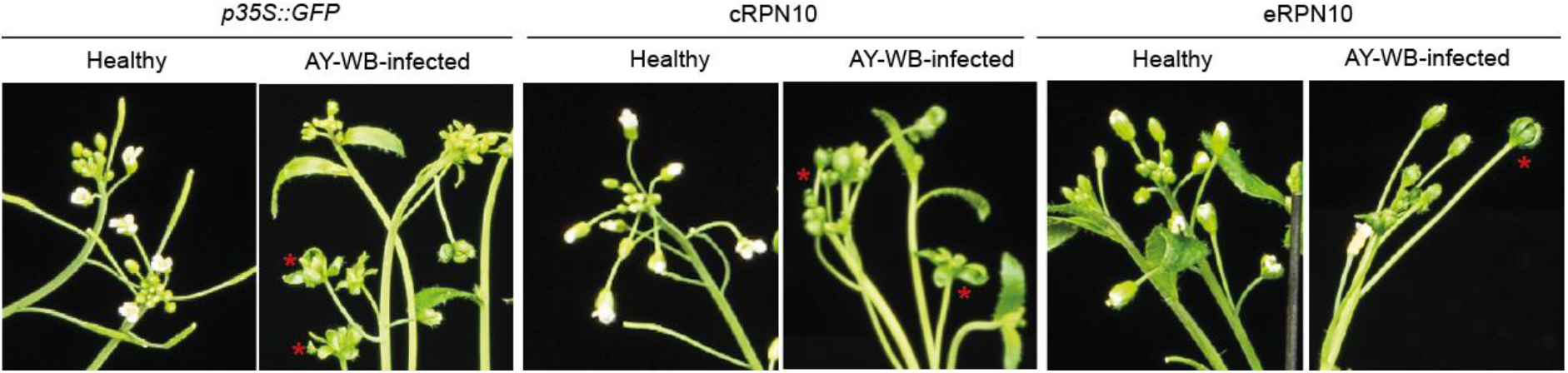
eRPN10 plants show phyllody symptoms during AY-WB infection. Typical flower morphology on healthy plants or infected plants is shown. Asterisks indicate leaf-like flowers on infected plants.

**Figure S10.**
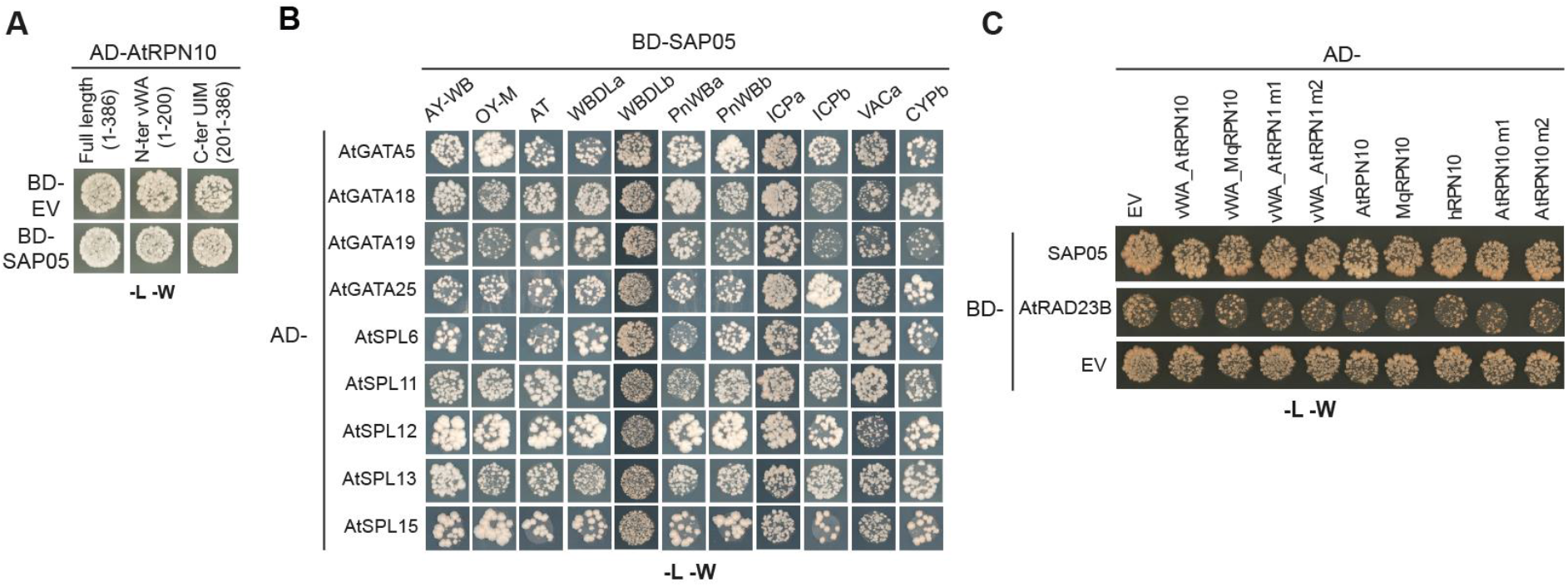
Yeast transformation controls for yeast two-hybrid assays in the study. (A-C) Yeast growth on medium lacking leucine and tryptophan, indicating presence of the AD and BD constructs in yeast two-hybrid assays for figures 2D (A), 4B (B) and 5B (C) in the main text.

**Table. S1.** Yeast two-hybrid screening of interacting proteins for the aster yellows witches’ broom phytoplasma effector SAP05 from an *A. thaliana* seedling cDNA library.

| Clone name | Gene name | Frame | Confidence<sup>a</sup> |
| --- | --- | --- | --- |
| pB27_A-64 | AtRPN10, AT4G38630 | In frame | A |
| pB27_A-134 | AtRPN10, AT4G38630 | In frame | A |
| pB27_A-51 | AtRPN10, AT4G38630 | In frame | A |
| pB27_A-142 | AtRPN10, AT4G38630 | In frame | A |
| pB27_A-109 | AtRPN10, AT4G38630 | In frame | A |
| pB27_A-40 | AtRPN10, AT4G38630 | In frame | A |
| pB27_A-49 | AtRPN10, AT4G38630 | In frame | A |
| pB27_A-54 | AtRPN10, AT4G38630 | In frame | A |
| pB27_A-93 | AtRPN10, AT4G38630 | In frame | A |
| pB27_A-13 | AtRPN10, AT4G38630 | In frame | A |
| pB27_A-33 | AtRPN10, AT4G38630 | In frame | A |
| pB27_A-3 | AtRPN10, AT4G38630 | In frame | A |
| pB27_A-68 | AtRPN10, AT4G38630 | In frame | A |
| pB27_A-125 | BME3 (GATA transcription factor 8), AT3G54810 | In frame | A |
| pB27_A-73 | BME3 (GATA transcription factor 8), AT3G54810 | In frame | A |
| pB27_A-55 | BME3 (GATA transcription factor 8), AT3G54810 | In frame | A |
| pB27_A-67 | BME3 (GATA transcription factor 8), AT3G54810 | In frame | A |
| pB27_A-36 | BME3 (GATA transcription factor 8), AT3G54810 | In frame | A |
| pB27_A-34 | BME3 (GATA transcription factor 8), AT3G54810 | In frame | A |
| pB27_A-101 | BME3 (GATA transcription factor 8), AT3G54810 | In frame | A |
| pB27_A-108 | BME3 (GATA transcription factor 8), AT3G54810 | In frame | A |
| pB27_A-75 | BME3 (GATA transcription factor 8), AT3G54810 | In frame | A |
| pB27_A-132 | BME3 (GATA transcription factor 8), AT3G54810 | In frame | A |
| pB27_A-89 | BME3 (GATA transcription factor 8), AT3G54810 | In frame | A |
| pB27_A-148 | BME3 (GATA transcription factor 8), AT3G54810 | In frame | A |
| pB27_A-92 | BME3 (GATA transcription factor 8), AT3G54810 | In frame | A |
| pB27_A-46 | BME3 (GATA transcription factor 8), AT3G54810 | In frame | A |
| pB27_A-1 | BME3 (GATA transcription factor 8), AT3G54810 | In frame | A |
| pB27_A-121 | BME3 (GATA transcription factor 8), AT3G54810 | In frame | A |
| pB27_A-70 | BME3 (GATA transcription factor 8), AT3G54810 | In frame | A |
| pB27_A-88 | BME3 (GATA transcription factor 8), AT3G54810 | In frame | A |
| pB27_A-128 | BME3 (GATA transcription factor 8), AT3G54810 | In frame | A |
| pB27_A-106 | BME3 (GATA transcription factor 8), AT3G54810 | In frame | A |
| pB27_A-35 | BME3 (GATA transcription factor 8), AT3G54810 | In frame | A |
| pB27_A-110 | BME3 (GATA transcription factor 8), AT3G54810 | In frame | A |
| pB27_A-12 | GNL/CGA1/AtGATA22 (GATA transcription factor 22), AT4G26150 | In frame | B |
| pB27_A-31 | GNL/CGA1/AtGATA22 (GATA transcription factor 22), AT4G26150 | In frame | B |
| pB27_A-53 | GNL/CGA1/AtGATA22 (GATA transcription factor 22), AT4G26150 | In frame | B |
| pB27_A-2 | GNL/CGA1/AtGATA22 (GATA transcription factor 22), AT4G26150 | In frame | B |
| pB27_A-149 | AtGATA1 (GATA transcription factor 1), AT3G24050 | In frame | B |
| pB27_A-47 | AtGATA1 (GATA transcription factor 1), AT3G24050 | In frame | B |
| pB27_A-57 | MNP/AtGATA18 (GATA transcription factor 18), AT3G50870 | In frame | B |
| pB27_A-86 | MNP/AtGATA18 (GATA transcription factor 18), AT3G50870 | In frame | B |
| pB27_A-5 | MNP/AtGATA18 (GATA transcription factor 18), AT3G50870 | In frame | B |
| pB27_A-83 | MNP/AtGATA18 (GATA transcription factor 18), AT3G50870 | In frame | B |
| pB27_A-133 | MNP/AtGATA18 (GATA transcription factor 18), AT3G50870 | In frame | B |
| pB27_A-17 | MNP/AtGATA18 (GATA transcription factor 18), AT3G50870 | In frame | B |
| pB27_A-143 | SPL14 (Squamosa promoter binding protein-like 14), AT1G20980 | In frame | D |
| pB27_A-22 | SPL14 (Squamosa promoter binding protein-like 14), AT1G20980 | N/A | D |
| pB27_A-72 | SPL13 (Squamosa promoter binding protein-like 13) AT5G50670 /AT5G50570 | In frame | D |
| pB27_A-120 | SPL13 (Squamosa promoter binding protein-like 13) AT5G50670 /AT5G50570 | In frame | D |
| pB27_A-116 | AtGATA9 (GATA transcription factor 9), AT4G32890 | In frame | B |
| pB27_A-85 | AtGATA9 (GATA transcription factor 9), AT4G32890 | In frame | B |
| pB27_A-16 | AtGATA9 (GATA transcription factor 9), AT4G32890 | In frame | B |
| pB27_A-61 | AtGATA5 (GATA transcription factor 5), AT5G66320 | N/A | N/A |
| pB27_A-140 | AtGATA27 (GATA transcription factor 27), AT5G47140 | In frame | B |
| pB27_A-98 | AtGATA27 (GATA transcription factor 27), AT5G47140 | In frame | B |
| pB27_A-19 | AtGATA27 (GATA transcription factor 27), AT5G47140 | In frame | B |
| pB27_A-117 | AtGATA27 (GATA transcription factor 27), AT5G47140 | In frame | B |
| pB27_A-20 | AtGATA27 (GATA transcription factor 27), AT5G47140 | N/A | N/A |
| pB27_A-77 | AtGATA27 (GATA transcription factor 27), AT5G47140 | N/A | N/A |
| pB27_A-26 | AtGATA7 (GATA transcription factor 7), AT4G36240 | In frame | D |
| pB27_A-87 | AtGATA19 (GATA transcription factor 19), AT4G36620 | In frame | D |
| pB27_A-69 | AtGATA19 (GATA transcription factor 19), AT4G36620 | In frame | D |
| pB27_A-91 | AtGATA19 (GATA transcription factor 19), AT4G36620 | In frame | D |
| pB27_A-147 | AtGATA19 (GATA transcription factor 19), AT4G36620 | In frame | D |
| pB27_A-119 | AtGATA19 (GATA transcription factor 19), AT4G36620 | In frame | D |
| pB27_A-138 | AtGATA19 (GATA transcription factor 19), AT4G36620 | In frame | D |
| pB27_A-104 | AtGATA4 (GATA transcription factor 4), AT3G60530 | In frame | D |
| pB27_A-44 | AtGATA4 (GATA transcription factor 4), AT3G60530 | In frame | D |

<sup>a</sup>Confidence level for interactions represented in the screen: A, very high confidence; B, high confidence; C, good confidence; D, moderate confidence; E, warning of non- specificity; F, experimentally proven artefacts; N/A, not attributed.

**Table. S2.** Summary of Y2H analysis of SAP05 interactions with GATA or SPL transcription factors.

| Gene name | Locus ID | Hybrigenics screening | *A. thaliana* TF library screening | Matchmaker gold system<sup>a</sup> Y2H | DUALhybrid Y2H system<sup>b</sup> |
| --- | --- | --- | --- | --- | --- |
| GATA1 | At3G24050 | + | - | + | NT |
| GATA2 | At2G45050 |  | - | + | NT |
| GATA3 | At4G34680 |  | - | + | NT |
| GATA4 | At3G60530 | + | - | + | NT |
| GATA5 | At5G66320 | + | - | + | NT |
| GATA6 | At3G51080 |  | - | + | NT |
| GATA7 | At4G36240 | + | - | + | NT |
| GATA8 (BME3) | At3G54810 | + | - | + | NT |
| GATA9 | At4G32890 | + | - | + | NT |
| GATA10 | At1G08000 |  | - | + | NT |
| GATA11 | At1G08010 |  | - | + | NT |
| GATA12 | At5G25830 |  | - | + | NT |
| GATA13 | At2G28340 |  | - | + | NT |
| GATA14 | At3G45170 |  | - | + | NT |
| GATA15 | At3G06740 |  | - | + | NT |
| GATA16 | At5G49300 |  | - | + | NT |
| GATA17 | At3G16870 |  | - | + | NT |
| GATA18 (HAN/MNP) | At3G50870 | + | + | + | NT |
| GATA19 | At4G36620 | + | + | + | NT |
| GATA20 | At2G18380 |  | + | + | NT |
| GATA21 (GNC) | At5G56860 |  |  | + | NT |
| GATA22 (GNL) | At4G26150 | + | - | - | NT |
| GATA23 | At5G26930 |  | - | + | NT |
| GATA24 (ZML1) | At3G21175 |  | - | + | NT |
| GATA25 (ZIM) | At4G24470 |  | + | + | NT |
| GATA26 | At4G17570 |  | + | NT | NT |
| GATA27 | At5G47140 | + | + | + | NT |
| GATA28 (ZML2) | At1G51600 |  | + | + | NT |
| SPL1 | At2G47070 |  | - | + | NT |
| SPL2 | At5G43270 |  | - | - | A.A. |
| SPL3 | At2G33810 |  | - | - | + |
| SPL4 | At1G53160 |  | - | - | + |
| SPL5 | At3G15270 |  | - | - | + |
| SPL6 | At1G69170 |  | + | + | NT |
| SPL7 | At5G18830 |  | - | - | - |
| SPL8 | At1G02065 |  | + | - | + |
| SPL9 | At2G42200 |  | + | - | A.A. |
| SPL10 | At1G27370 |  | - | - | A.A. |
| SPL11 | At1G27360 |  | - | + | NT |
| SPL12 | At3G60030 |  | - | + | NT |
| SPL13A | At5G50570 |  | + | + | NT |
| SPL13B | At5G50670 | + | + | + | NT |
| SPL14 | At1G20980 | + | - | + | - |
| SPL15 | At3G57920 |  | + | + | NT |
| SPL16 | At1G76580 |  | - | NT | NT |

<sup>a</sup> '+' indicates yeast growth; '-' indicates no yeast growth; NT, non-tested; A.A., autoactivation.
<sup>a</sup>SAP05(33-135) was cloned into the pGBKT7 plasmid, and GATA or SPL transcription factors into the pGADT7 plasmid. Yeast strain AH109 was used for transformation and growth analysis.
<sup>b</sup>SAP05(33-135) was cloned into the pGADHA plasmid, and GATA or SPL transcription factors into the pLEXA-C plasmid as N-terminal fusions. Yeast strain NMY51 was used for transformation and growth analysis.

**Table. S4.**
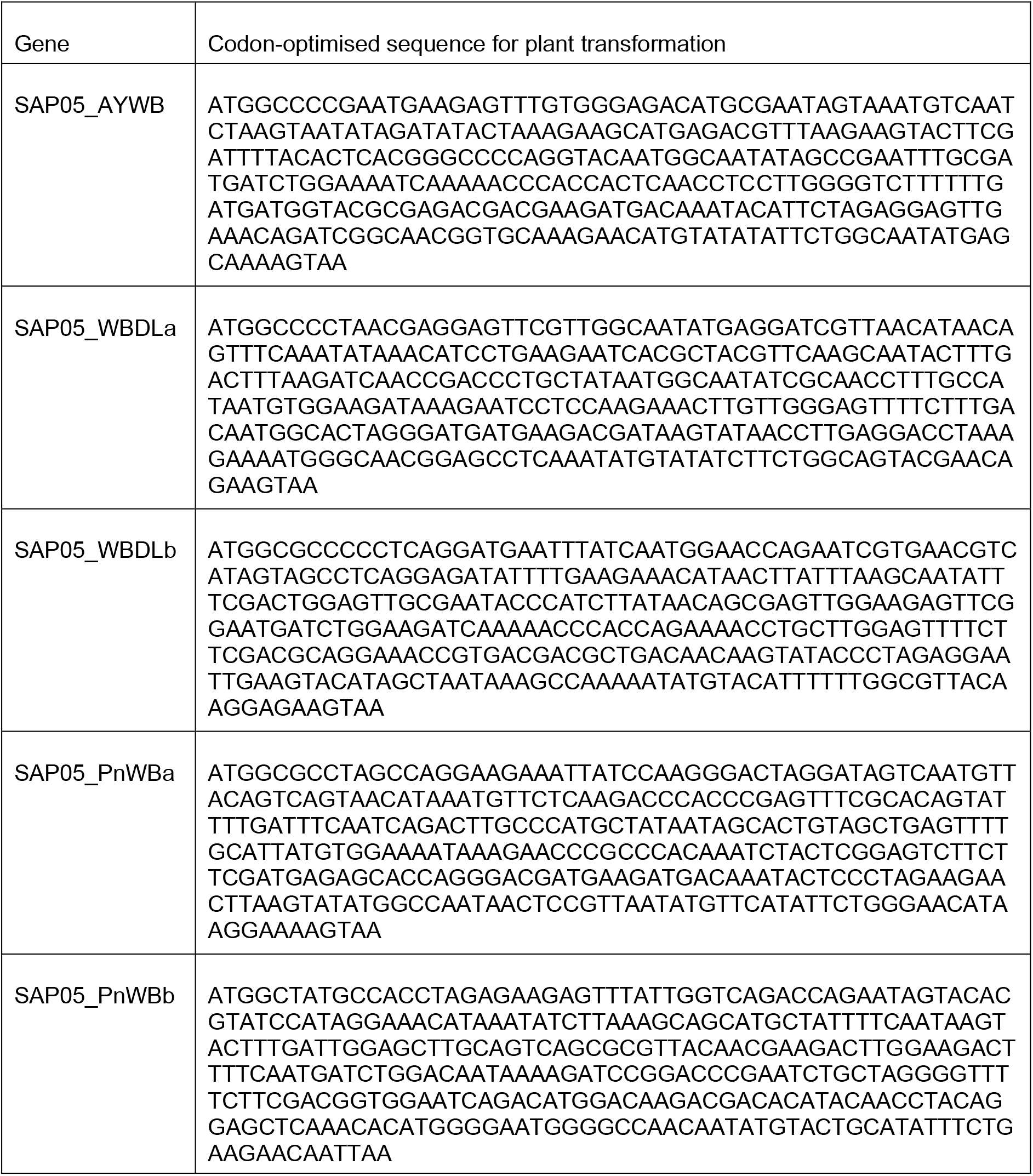
Sequences of codon optimized SAP05 genes for generating transgenic plants.

**Table. S5.** Plasmids used in the study.

| Plasmid name | Vector | Purpose |
| --- | --- | --- |
| pB7WG2-SAP05_AYWB | pB7WG2 | Generating *p35S::SAP05_AYWB* plants and transient expression in protoplasts |
| pHW59-SAP05_AYWB | pHW59 | Generating *pATSUC2::SAP05* plants |
| pB7WG2-SAP05_WBDLa | pB7WG2 | Generating *p35S::SAP05_WBDLa* plants and transient expression in protoplasts |
| pB7WG2-SAP05_WBDLb | pB7WG2 | Generating *p35S::SAP05_WBDLb* plants and transient expression in protoplasts |
| pB7WG2-SAP05_PnWBa | pB7WG2 | Generating *p35S::SAP05_PnWBa* plants and transient expression in protoplasts |
| pB7WG2-SAP05_PnWBb | pB7WG2 | Generating *p35S::SAP05_PnWBb* plants and transient expression in protoplasts |
| pB7WG2-SAP05_AT | pB7WG2 | Transient expression in protoplasts |
| pB7WG2-SAP05_VACa | pB7WG2 | Transient expression in protoplasts |
| pB7WG2-GFP | pB7WG2 | Generating GFP control plants and transient expression in protoplasts |
| pB7WG2-AtRPN10 | pB7WG2 | Transient expression in protoplasts |
| pB7WG2-MqRPN10 | pB7WG2 | Transient expression in protoplasts |
| pB7WG2-hRPN10 | pB7WG2 | Transient expression in protoplasts |
| pB7WG2-AtRPN10 m1 | pB7WG2 | Transient expression in protoplasts |
| pB7WG2-AtRPN10 m2 | pB7WG2 | Transient expression in protoplasts |
| pBI121-*pAtRPN10::AtRPN10* | pBI121 | Generating cRPN10 plants |
| pBI121-*pAtRPN10::AtRPN10 m1* | pBI121 | Generating eRPN10 plants |
| pGADT7 | pGADT7 | Y2H Empty vector (Matchmaker system) |
| pGBKT7 | pGABK7 | Y2H Empty vector (Matchmaker system) |
| pGBKT7-SAP05_AYWB | pGBKT7 | Y2H analysis (Matchmaker system) |
| pGBKT7-SAP05_WBDLa | pGBKT7 | Y2H analysis (Matchmaker system) |
| pGBKT7-SAP05_WBDLb | pGBKT7 | Y2H analysis (Matchmaker system) |
| pGBKT7-SAP05_PnWBa | pGBKT7 | Y2H analysis (Matchmaker system) |
| pGBKT7-SAP05_PnWBb | pGBKT7 | Y2H analysis (Matchmaker system) |
| pGBKT7-SAP05_OY-M | pGBKT7 | Y2H analysis (Matchmaker system) |
| pGBKT7-SAP05_AT | pGBKT7 | Y2H analysis (Matchmaker system) |
| pGBKT7-SAP05_ICPa | pGBKT7 | Y2H analysis (Matchmaker system) |
| pGBKT7-SAP05_ICPb | pGBKT7 | Y2H analysis (Matchmaker system) |
| pGBKT7-SAP05_VACa | pGBKT7 | Y2H analysis (Matchmaker system) |
| pGBKT7-SAP05_CYPb | pGBKT7 | Y2H analysis (Matchmaker system) |
| pGBKT7-SAP05_AtRAD23B | pGBKT7 | Y2H analysis (Matchmaker system) |
| pGADT7-AtRPN10 | pGADT7 | Y2H analysis (Matchmaker system) |
| pGADT7-AtRPN10_vWA | pGADT7 | Y2H analysis (Matchmaker system) |
| pGADT7-AtRPN10_UIM | pGADT7 | Y2H analysis (Matchmaker system) |
| pGADT7-MqRPN10 | pGADT7 | Y2H analysis (Matchmaker system) |
| pGADT7-hRPN10 | pGADT7 | Y2H analysis (Matchmaker system) |
| pGADT7-AtRPN10 m1 | pGADT7 | Y2H analysis (Matchmaker system) |
| pGADT7-AtRPN10 m2 | pGADT7 | Y2H analysis (Matchmaker system) |
| pGADT7-vWA_AtRPN10 | pGADT7 | Y2H analysis (Matchmaker system) |
| pGADT7-vWA_MqRPN10 | pGADT7 | Y2H analysis (Matchmaker system) |
| pGADT7-vWA_AtRPN10 m1 | pGADT7 | Y2H analysis (Matchmaker system) |
| pGADT7-vWA_AtRPN10 m2 | pGADT7 | Y2H analysis (Matchmaker system) |
| pGADT7-GATA1 | pGADT7 | Y2H analysis (Matchmaker system) |
| pGADT7-GATA2 | pGADT7 | Y2H analysis (Matchmaker system) |
| pGADT7-GATA3 | pGADT7 | Y2H analysis (Matchmaker system) |
| pGADT7-GATA4 | pGADT7 | Y2H analysis (Matchmaker system) |
| pGADT7-GATA5 | pGADT7 | Y2H analysis (Matchmaker system) |
| pGADT7-GATA6 | pGADT7 | Y2H analysis (Matchmaker system) |
| pGADT7-GATA7 | pGADT7 | Y2H analysis (Matchmaker system) |
| pGADT7-GATA8 | pGADT7 | Y2H analysis (Matchmaker system) |
| pGADT7-GATA9 | pGADT7 | Y2H analysis (Matchmaker system) |
| pGADT7-GATA10 | pGADT7 | Y2H analysis (Matchmaker system) |
| pGADT7-GATA11 | pGADT7 | Y2H analysis (Matchmaker system) |
| pGADT7-GATA12 | pGADT7 | Y2H analysis (Matchmaker system) |
| pGADT7-GATA14 | pGADT7 | Y2H analysis (Matchmaker system) |
| pGADT7-GATA15 | pGADT7 | Y2H analysis (Matchmaker system) |
| pGADT7-GATA16 | pGADT7 | Y2H analysis (Matchmaker system) |
| pGADT7-GATA17 | pGADT7 | Y2H analysis (Matchmaker system) |
| pGADT7-GATA18 | pGADT7 | Y2H analysis (Matchmaker system) |
| pGADT7-GATA19 | pGADT7 | Y2H analysis (Matchmaker system) |
| pGADT7-GATA20 | pGADT7 | Y2H analysis (Matchmaker system) |
| pGADT7-GATA21 | pGADT7 | Y2H analysis (Matchmaker system) |
| pGADT7-GATA22 | pGADT7 | Y2H analysis (Matchmaker system) |
| pGADT7-GATA24 | pGADT7 | Y2H analysis (Matchmaker system) |
| pGADT7-GATA25 | pGADT7 | Y2H analysis (Matchmaker system) |
| pGADT7-GATA27 | pGADT7 | Y2H analysis (Matchmaker system) |
| pGADT7-MqGATA ZnF1 | pGADT7 | Y2H analysis (Matchmaker system) |
| pGADT7-MqGATA ZnF2 | pGADT7 | Y2H analysis (Matchmaker system) |
| pGADT7-MqGATA ZnF3 | pGADT7 | Y2H analysis (Matchmaker system) |
| pGADT7-MqGATA ZnF4 | pGADT7 | Y2H analysis (Matchmaker system) |
| pGADT7-MqGATA ZnF5 | pGADT7 | Y2H analysis (Matchmaker system) |
| pGADT7-MqGATA ZnF6 | pGADT7 | Y2H analysis (Matchmaker system) |
| pGADT7-SPL1 | pGADT7 | Y2H analysis (Matchmaker system) |
| pGADT7-SPL2 | pGADT7 | Y2H analysis (Matchmaker system) |
| pGADT7-SPL3 | pGADT7 | Y2H analysis (Matchmaker system) |
| pGADT7-SPL4 | pGADT7 | Y2H analysis (Matchmaker system) |
| pGADT7-SPL5 | pGADT7 | Y2H analysis (Matchmaker system) |
| pGADT7-SPL6 | pGADT7 | Y2H analysis (Matchmaker system) |
| pGADT7-SPL7 | pGADT7 | Y2H analysis (Matchmaker system) |
| pGADT7-SPL8 | pGADT7 | Y2H analysis (Matchmaker system) |
| pGADT7-SPL9 | pGADT7 | Y2H analysis (Matchmaker system) |
| pGADT7-SPL10 | pGADT7 | Y2H analysis (Matchmaker system) |
| pGADT7-SPL11 | pGADT7 | Y2H analysis (Matchmaker system) |
| pGADT7-SPL12 | pGADT7 | Y2H analysis (Matchmaker system) |
| pGADT7-SPL13 | pGADT7 | Y2H analysis (Matchmaker system) |
| pGADT7-SPL14 | pGADT7 | Y2H analysis (Matchmaker system) |
| pGADT7-SPL15 | pGADT7 | Y2H analysis (Matchmaker system) |
| PGADT7-ZnF_GATA18 | pGADT7 | GATA domain of GATA15 for Y2H |
| PGADT7-ZnF_SPL13 | pGADT7 | SBP domain of SPL13 for Y2H |
| PGADT7-ZnF_SPL15 | pGADT7 | SBP domain of SPL15 for Y2H |
| pGADHA | pGADHA | Y2H Empty vector (DUALhybrid system) |
| pLEXA-C | pLEXA-C | Y2H Empty vector (DUALhybrid s system) |
| pGADHA-SAP05 | pGADHA | Y2H analysis (Matchmaker system) |
| pLEXA-C-SPL2 | pLEXA-C | Y2H analysis (Matchmaker system) |
| pLEXA-C-SPL3 | pLEXA-C | Y2H analysis (Matchmaker system) |
| pLEXA-C-SPL4 | pLEXA-C | Y2H analysis (Matchmaker system) |
| pLEXA-C-SPL5 | pLEXA-C | Y2H analysis (Matchmaker system) |
| pLEXA-C-SPL7 | pLEXA-C | Y2H analysis (Matchmaker system) |
| pLEXA-C-SPL8 | pLEXA-C | Y2H analysis (Matchmaker system) |
| pLEXA-C-SPL9 | pLEXA-C | Y2H analysis (Matchmaker system) |
| pLEXA-C-SPL10 | pLEXA-C | Y2H analysis (Matchmaker system) |
| pLEXA-C-SPL14 | pLEXA-C | Y2H analysis (Matchmaker system) |
| pUGW15-GATA19 | pUGW15 | Protoplast expression with a N-ter HA tag |
| pUGW15-GATA18 | pUGW15 | Protoplast expression with a N-ter HA tag |
| pUGW15-GATA18 (K->R) | pUGW15 | Protoplast expression with a N-ter HA tag |
| pUGW15-GATA19 (K->R) | pUGW15 | Protoplast expression with a N-ter HA tag |
| pUGW15-SPL5 | pUGW15 | Protoplast expression with a N-ter HA tag |
| pUGW15-rSPL9 | pUGW15 | Protoplast expression with a N-ter HA tag |
| pUGW15-rSPL11 | pUGW15 | Protoplast expression with a N-ter HA tag |
| pUGW15-rSPL13 | pUGW15 | Protoplast expression with a N-ter HA tag |
| pUGW15-rSPL15 | pUGW15 | Protoplast expression with a N-ter HA tag |
| pUGW15-ZnF_GATA18 | pUGW15 | Protoplast expression with a N-ter HA tag |
| pUGW15-ZnF_SPL13 | pUGW15 | Protoplast expression with a N-ter HA tag |
| pOPINF-SAP05 | pOPINF | 6XHis-3C-SAP05 expression in *E.coli* |
| pOPINF-ZnF_SPL5 | pOPINF | 6XHis-3C-ZnF_SPL5 expression in *E.coli* |
| pOPINA-SUMO-vWA | pOPINA | SUMO-vWA expression in *E.coli* |
| pOPINA-SUMO-evWA | pOPINA | SUMO-evWA expression in *E.coli* |
| pOPINS3C-vWA | pOPINS3C | 6XHis-SUMO-3C-vWA expression in *E.coli* |
| pOPINS3C-evWA | pOPINS3C | 6XHis-SUMO-3C-evWA expression in *E.coli* |

## Notes

### Competing Interest Statement

The authors have declared no competing interest.

## References

Abramovitch, R.B., Janjusevic, R., Stebbins, C.E., and Martin, G.B. (2006). Type III effector AvrPtoB requires intrinsic E3 ubiquitin ligase activity to suppress plant cell death and immunity. Proc Natl Acad Sci U S A 103, 2851–2856.

Al-Subhi, A.M., Al-Sadi, A.M., Al-Yahyai, R., Chen, Y., Mathers, T., Orlovskis, Z., Moro, G., Mugford, S., Al-Hashmi, K., and Hogenhout, S. (2020). Witches’ broom disease of lime contributes to phytoplasma epidemics and attracts insect vectors. Plant Dis. doi: 10.1094/PDIS-10-20-2112-RE. Online ahead of print.

Arashida, R., Kakizawa, S., Ishii, Y., Hoshi, A., Jung, H.Y., Kagiwada, S., Yamaji, Y., Oshima, K., and Namba, S. (2008). Cloning and characterization of the antigenic membrane protein (Amp) gene and in situ detection of Amp from malformed flowers infected with Japanese hydrangea phyllody phytoplasma. Phytopathology 98, 769–775.

Ashida, H., Kim, M., and Sasakawa, C. (2014). Exploitation of the host ubiquitin system by human bacterial pathogens. Nature Reviews Microbiology 12, 399–413.

Bai, X., Correa, V.R., Toruno, T.Y., Ammar el, D., Kamoun, S., and Hogenhout, S.A. (2009). AY-WB phytoplasma secretes a protein that targets plant cell nuclei. Mol Plant Microbe Interact 22, 18–30.

Bai, X., Zhang, J., Ewing, A., Miller, S.A., Jancso Radek, A., Shevchenko, D.V., Tsukerman, K., Walunas, T., Lapidus, A., Campbell, J.W., et al. (2006). Living with genome instability: the adaptation of phytoplasmas to diverse environments of their insect and plant hosts. J Bacteriol 188, 3682–3696.

Banfield, M.J. (2015). Perturbation of host ubiquitin systems by plant pathogen/pest effector proteins. Cell Microbiol 17, 18–25.

Beanland, L., Hoy, C.W., Miller, S.A., and Nault, L.R. (2000). Influence of Aster Yellows Phytoplasma on the Fitness of Aster Leafhopper (Homoptera: Cicadellidae). Ann Entomol Soc Am 93, 271–276.

Berrow, N.S., Alderton, D., Sainsbury, S., Nettleship, J., Assenberg, R., Rahman, N., Stuart, D.I., and Owens, R.J. (2007). A versatile ligation-independent cloning method suitable for high-throughput expression screening applications. Nucleic Acids Res 35, e45.

CABI (2021). Invasive Species Compendium. Wallingford, UK: CAB international. www.cabi.org/isc.

Cano, L.M., Raffaele, S., Haugen, R.H., Saunders, D.G., Leonelli, L., MacLean, D., Hogenhout, S.A., and Kamoun, S. (2013). Major transcriptome reprogramming underlies floral mimicry induced by the rust fungus *Puccinia monoica* in Boechera stricta. PLoS One 8, e75293.

Chamberlain, P.P., and Hamann, L.G. (2019). Development of targeted protein degradation therapeutics. Nat Chem Biol 15, 937–944.

Chang, S.H., Tan, C.M., Wu, C.T., Lin, T.H., Jiang, S.Y., Liu, R.C., Tsai, M.C., Su, L.W., and Yang, J.Y. (2018). Alterations of plant architecture and phase transition by the phytoplasma virulence factor SAP11. J Exp Bot 69, 5389–5401.

Chen, K., Wang, Y., Zhang, R., Zhang, H., and Gao, C. (2019). CRISPR/Cas Genome Editing and Precision Plant Breeding in Agriculture. Annu Rev Plant Biol 70, 667–697.

Chung, W.C., Chen, L.L., Lo, W.S., Lin, C.P., and Kuo, C.H. (2013). Comparative analysis of the peanut witches’-broom phytoplasma genome reveals horizontal transfer of potential mobile units and effectors. PLoS One 8, e62770.

Cui, J., Yao, Q., Li, S., Ding, X., Lu, Q., Mao, H., Liu, L., Zheng, N., Chen, S., and Shao, F. (2010). Glutamine deamidation and dysfunction of ubiquitin/NEDD8 induced by a bacterial effector family. Science 329, 1215–1218.

Dawkins, R. (1982). The Extended Phenotype: The Long Reach of the Gene (Oxford: Oxford University Press).

Ding, L., Yan, S., Jiang, L., Liu, M., Zhang, J., Zhao, J., Zhao, W., Han, Y.Y., Wang, Q., and Zhang, X. (2015). HANABA TARANU regulates the shoot apical meristem and leaf development in cucumber (*Cucumis sativus* L.). J Exp Bot 66, 7075–7087.

Doi, Y., Teranaka, M., Yora, K., and Asuyama, H. (1967). Mycoplasma- or PLT Group-like Microorganisms Found in the Phloem Elements of Plants Infected with Mulberry Dwarf, Potato Witches’ Broom, Aster Yellows, or Paulownia Witches’ Broom. Japanese Journal of Phytopathology 33, 259–266.

Drurey, C., Mathers, T.C., Prince, D.C., Wilson, C., Caceres-Moreno, C., Mugford, S.T., and Hogenhout, S.A. (2019). Chemosensory proteins in the CSP4 clade evolved as plant immunity suppressors before two suborders of plant-feeding hemipteran insects diverged. bioRxiv 173278; doi: https://doi.org/10.1101/173278.

Du Toit, A. (2014). Bacterial pathogenicity: Phytoplasma converts plants into zombies. Nat Rev Microbiol 12, 393.

EPPO (2021). EPPO Global Database (available online). https://gd.eppo.int.

Farmer, L.M., Book, A.J., Lee, K.H., Lin, Y.L., Fu, H., and Vierstra, R.D. (2010). The RAD23 family provides an essential connection between the 26S proteasome and ubiquitylated proteins in Arabidopsis. Plant Cell 22, 124–142.

Franco-Zorrilla, J.M., Valli, A., Todesco, M., Mateos, I., Puga, M.I., Rubio-Somoza, I., Leyva, A., Weigel, D., Garcia, J.A., and Paz-Ares, J. (2007). Target mimicry provides a new mechanism for regulation of microRNA activity. Nat Genet 39, 1033–1037.

Frost, K.E., Esker, P.D., Van Haren, R., Kotolski, L., and Groves, R.L. (2013a). Factors influencing aster leafhopper (Hemiptera: Cicadellidae) abundance and aster yellows phytoplasma infectivity in Wisconsin carrot fields. Environ Entomol 42, 477–490.

Frost, K.E., Esker, P.D., Van Haren, R., Kotolski, L., and Groves, R.L. (2013b). Seasonal patterns of aster leafhopper (Hemiptera: Cicadellidae) abundance and aster yellows phytoplasma infectivity in Wisconsin carrot fields. Environ Entomol 42, 491–502.

Frost, K.E., Willis, D.K., and Groves, R.L. (2011). Detection and variability of aster yellows phytoplasma titer in its insect vector, *Macrosteles quadrilineatus* (Hemiptera: Cicadellidae). J Econ Entomol 104, 1800–1815.

Fu, H., Sadis, S., Rubin, D.M., Glickman, M., van Nocker, S., Finley, D., and Vierstra, R.D. (1998). Multiubiquitin chain binding and protein degradation are mediated by distinct domains within the 26 S proteasome subunit Mcb1. J Biol Chem 273, 1970–1981.

Groll, M., Schellenberg, B., Bachmann, A.S., Archer, C.R., Huber, R., Powell, T.K., Lindow, S., Kaiser, M., and Dudler, R. (2008). A plant pathogen virulence factor inhibits the eukaryotic proteasome by a novel mechanism. Nature 452, 755–758.

Herbison, R., Lagrue, C., and Poulin, R. (2018). The missing link in parasite manipulation of host behaviour. Parasit Vectors 11, 222.

Hogenhout, S.A., Oshima, K., Ammar el, D., Kakizawa, S., Kingdom, H.N., and Namba, S. (2008). Phytoplasmas: bacteria that manipulate plants and insects. Mol Plant Pathol 9, 403–423.

Hoshi, A., Oshima, K., Kakizawa, S., Ishii, Y., Ozeki, J., Hashimoto, M., Komatsu, K., Kagiwada, S., Yamaji, Y., and Namba, S. (2009). A unique virulence factor for proliferation and dwarfism in plants identified from a phytopathogenic bacterium. Proc Natl Acad Sci U S A 106, 6416–6421.

Huang, W., Reyes-Caldas, P., Mann, M., Seifbarghi, S., Kahn, A., Almeida, R.P.P., Beven, L., Heck, M., Hogenhout, S.A., and Coaker, G. (2020). Bacterial Vector-Borne Plant Diseases: Unanswered Questions and Future Directions. Molecular plant 13, 1379–1393.

Hudson, D., Guevara, D.R., Hand, A.J., Xu, Z., Hao, L., Chen, X., Zhu, T., Bi, Y.M., and Rothstein, S.J. (2013). Rice cytokinin GATA transcription Factor1 regulates chloroplast development and plant architecture. Plant Physiol 162, 132–144.

Hughes, D.P., and Libersat, F. (2019). Parasite manipulation of host behavior. Curr Biol 29, R45–r47.

Huijser, P., and Schmid, M. (2011). The control of developmental phase transitions in plants. Development 138, 4117–4129.

IRPCM (2004). ’*Candidatus* Phytoplasma’, a taxon for the wall-less, non-helical prokaryotes that colonize plant phloem and insects. Int J Syst Evol Microbiol 54, 1243–1255.

Iwabuchi, N., Kitazawa, Y., Maejima, K., Koinuma, H., Miyazaki, A., Matsumoto, O., Suzuki, T., Nijo, T., Oshima, K., Namba, S., et al. (2020). Functional variation in phyllogen, a phyllody-inducing phytoplasma effector family, attributable to a single amino acid polymorphism. Mol Plant Pathol 21, 1322–1336.

Johnson, H.J., and Koshy, A.A. (2020). Latent Toxoplasmosis Effects on Rodents and Humans: How Much is Real and How Much is Media Hype? mBio 11.

Johnson, P.T., Lunde, K.B., Ritchie, E.G., and Launer, A.E. (1999). The effect of trematode infection on amphibian limb development and survivorship. Science 284, 802–804.

Kakizawa, S., Oshima, K., Nishigawa, H., Jung, H.Y., Wei, W., Suzuki, S., Tanaka, M., Miyata, S.I., Ugaki, M., and Namba, S. (2004). Secretion of immunodominant membrane protein from onion yellows phytoplasma through the Sec protein-translocation system in *Escherichia coli*. Microbiology 150, 135–142.

Kitazawa, Y., Iwabuchi, N., Himeno, M., Sasano, M., Koinuma, H., Nijo, T., Tomomitsu, T., Yoshida, T., Okano, Y., Yoshikawa, N., et al. (2017). Phytoplasma-conserved phyllogen proteins induce phyllody across the Plantae by degrading floral MADS domain proteins. J Exp Bot 68, 2799–2811.

Koinuma, H., Maejima, K., Tokuda, R., Kitazawa, Y., Nijo, T., Wei, W., Kumita, K., Miyazaki, A., Namba, S., and Yamaji, Y. (2020). Spatiotemporal dynamics and quantitative analysis of phytoplasmas in insect vectors. Sci Rep 10, 4291.

Le Fevre, R., Evangelisti, E., Rey, T., and Schornack, S. (2015). Modulation of host cell biology by plant pathogenic microbes. Annu Rev Cell Dev Biol 31, 201–229.

Lee, I.M., Davis, R.E., and Gundersen-Rindal, D.E. (2000). Phytoplasma: phytopathogenic mollicutes. Annu Rev Microbiol 54, 221–255.

Lin, Y.H., and Machner, M.P. (2017). Exploitation of the host cell ubiquitin machinery by microbial effector proteins. J Cell Sci 130, 1985–1996.

Lin, Y.L., Sung, S.C., Tsai, H.L., Yu, T.T., Radjacommare, R., Usharani, R., Fatimababy, A.S., Lin, H.Y., Wang, Y.Y., and Fu, H. (2011). The defective proteasome but not substrate recognition function is responsible for the null phenotypes of the Arabidopsis proteasome subunit RPN10. Plant Cell 23, 2754–2773.

Lu, Y.T., Li, M.Y., Cheng, K.T., Tan, C.M., Su, L.W., Lin, W.Y., Shih, H.T., Chiou, T.J., and Yang, J.Y. (2014). Transgenic plants that express the phytoplasma effector SAP11 show altered phosphate starvation and defense responses. Plant Physiol 164, 1456–1469.

MacLean, A.M., Orlovskis, Z., Kowitwanich, K., Zdziarska, A.M., Angenent, G.C., Immink, R.G.H., and Hogenhout, S.A. (2014). Phytoplasma Effector SAP54 Hijacks Plant Reproduction by Degrading MADS-box Proteins and Promotes Insect Colonization in a RAD23-Dependent Manner. PLoS Biol 12.

MacLean, A.M., Sugio, A., Makarova, O.V., Findlay, K.C., Grieve, V.M., Toth, R., Nicolaisen, M., and Hogenhout, S.A. (2011). Phytoplasma effector SAP54 induces indeterminate leaf-like flower development in Arabidopsis plants. Plant Physiol 157, 831–841.

Maejima, K., Iwai, R., Himeno, M., Komatsu, K., Kitazawa, Y., Fujita, N., Ishikawa, K., Fukuoka, M., Minato, N., Yamaji, Y., et al. (2014). Recognition of floral homeotic MADS domain transcription factors by a phytoplasmal effector, phyllogen, induces phyllody. Plant J 78, 541–554.

Malembic-Maher, S., Desque, D., Khalil, D., Salar, P., Bergey, B., Danet, J.L., Duret, S., Dubrana-Ourabah, M.P., Beven, L., Ember, I., et al. (2020). When a Palearctic bacterium meets a Nearctic insect vector: Genetic and ecological insights into the emergence of the grapevine Flavescence doree epidemics in Europe. PLoS Pathog 16, e1007967.

Marshall, R.S., Li, F., Gemperline, D.C., Book, A.J., and Vierstra, R.D. (2015). Autophagic Degradation of the 26S Proteasome Is Mediated by the Dual ATG8/Ubiquitin Receptor RPN10 in Arabidopsis. Mol Cell 58, 1053–1066.

Mathieu, J., Warthmann, N., Kuttner, F., and Schmid, M. (2007). Export of FT protein from phloem companion cells is sufficient for floral induction in Arabidopsis. Curr Biol 17, 1055–1060.

Mayor, T., Graumann, J., Bryan, J., MacCoss, M.J., and Deshaies, R.J. (2007). Quantitative profiling of ubiquitylated proteins reveals proteasome substrates and the substrate repertoire influenced by the Rpn10 receptor pathway. Mol Cell Proteomics 6, 1885–1895.

Musetti, R., Buxa, S.V., De Marco, F., Loschi, A., Polizzotto, R., Kogel, K.H., and van Bel, A.J. (2013). Phytoplasma-triggered Ca(2+) influx is involved in sieve-tube blockage. Mol Plant Microbe Interact 26, 379–386.

Nault (1990). Evolution of an insect pest: maize and the corn leafhopper, a case study. Maydica 35, 165–175.

Orlovskis, Z., and Hogenhout, S.A. (2016). A Bacterial Parasite Effector Mediates Insect Vector Attraction in Host Plants Independently of Developmental Changes. Frontiers in plant science 7, 885.

Oshima, K., Ishii, Y., Kakizawa, S., Sugawara, K., Neriya, Y., Himeno, M., Minato, N., Miura, C., Shiraishi, T., Yamaji, Y., et al. (2011). Dramatic transcriptional changes in an intracellular parasite enable host switching between plant and insect. PLoS One 6, e23242.

Oshima, K., Kakizawa, S., Nishigawa, H., Jung, H.Y., Wei, W., Suzuki, S., Arashida, R., Nakata, D., Miyata, S., Ugaki, M., et al. (2004). Reductive evolution suggested from the complete genome sequence of a plant-pathogenic phytoplasma. Nat Genet 36, 27–29.

Pecher, P., Moro, G., Canale, M.C., Capdevielle, S., Singh, A., MacLean, A., Sugio, A., Kuo, C.H., Lopes, J.R.S., and Hogenhout, S.A. (2019). Phytoplasma SAP11 effector destabilization of TCP transcription factors differentially impact development and defence of Arabidopsis versus maize. PLoS Pathog 15, e1008035.

Pruneda-Paz, J.L., Breton, G., Nagel, D.H., Kang, S.E., Bonaldi, K., Doherty, C.J., Ravelo, S., Galli, M., Ecker, J.R., and Kay, S.A. (2014). A genome-scale resource for the functional characterization of Arabidopsis transcription factors. Cell Rep 8, 622–632.

Qiu, J.Z., Sheedlo, M.J., Yu, K.W., Tan, Y.H., Nakayasu, E.S., Das, C., Liu, X.Y., and Luo, Z.Q. (2016). Ubiquitination independent of E1 and E2 enzymes by bacterial effectors. Nature 533, 120–124.

Ranftl, Q.L., Bastakis, E., Klermund, C., and Schwechheimer, C. (2016). LLM-Domain Containing B-GATA Factors Control Different Aspects of Cytokinin-Regulated Development in Arabidopsis thaliana. Plant Physiol 170, 2295–2311.

Richter, R., Bastakis, E., and Schwechheimer, C. (2013). Cross-repressive interactions between SOC1 and the GATAs GNC and GNL/CGA1 in the control of greening, cold tolerance, and flowering time in Arabidopsis. Plant Physiol 162, 1992–2004.

Richter, R., Behringer, C., Muller, I.K., and Schwechheimer, C. (2010). The GATA-type transcription factors GNC and GNL/CGA1 repress gibberellin signaling downstream from DELLA proteins and PHYTOCHROME-INTERACTING FACTORS. Genes Dev 24, 2093–2104.

Riedinger, C., Boehringer, J., Trempe, J.F., Lowe, E.D., Brown, N.R., Gehring, K., Noble, M.E., Gordon, C., and Endicott, J.A. (2010). Structure of Rpn10 and its interactions with polyubiquitin chains and the proteasome subunit Rpn12. J Biol Chem 285, 33992–34003.

Rothbauer, U., Zolghadr, K., Tillib, S., Nowak, D., Schermelleh, L., Gahl, A., Backmann, N., Conrath, K., Muyldermans, S., Cardoso, M.C., et al. (2006). Targeting and tracing antigens in live cells with fluorescent nanobodies. Nat Methods 3, 887–889.

Rumpler, F., Gramzow, L., Theissen, G., and Melzer, R. (2015). Did Convergent Protein Evolution Enable Phytoplasmas to Generate ‘Zombie Plants’? Trends Plant Sci 20, 798–806.

Samad, A.F.A., Sajad, M., Nazaruddin, N., Fauzi, I.A., Murad, A.M.A., Zainal, Z., and Ismail, I. (2017). MicroRNA and Transcription Factor: Key Players in Plant Regulatory Network. Frontiers in plant science 8, 565.

Sanada, T., Kim, M., Mimuro, H., Suzuki, M., Ogawa, M., Oyama, A., Ashida, H., Kobayashi, T., Koyama, T., Nagai, S., et al. (2012). The *Shigella flexneri* effector OspI deamidates UBC13 to dampen the inflammatory response. Nature 483, 623–626.

Schapira, M., Calabrese, M.F., Bullock, A.N., and Crews, C.M. (2019). Targeted protein degradation: expanding the toolbox. Nat Rev Drug Discov. 18, 949–963.

Shapira, M., Hamlin, B.J., Rong, J., Chen, K., Ronen, M., and Tan, M.W. (2006). A conserved role for a GATA transcription factor in regulating epithelial innate immune responses. Proc Natl Acad Sci U S A 103, 14086–14091.

Smalle, J., Kurepa, J., Yang, P., Emborg, T.J., Babiychuk, E., Kushnir, S., and Vierstra, R.D. (2003). The pleiotropic role of the 26S proteasome subunit RPN10 in Arabidopsis growth and development supports a substrate-specific function in abscisic acid signaling. Plant Cell 15, 965–980.

Sugio, A., Kingdom, H.N., MacLean, A.M., Grieve, V.M., and Hogenhout, S.A. (2011a). Phytoplasma protein effector SAP11 enhances insect vector reproduction by manipulating plant development and defense hormone biosynthesis. Proc Natl Acad Sci U S A 108, E1254–1263.

Sugio, A., MacLean, A.M., Kingdom, H.N., Grieve, V.M., Manimekalai, R., and Hogenhout, S.A. (2011b). Diverse targets of phytoplasma effectors: from plant development to defense against insects. Annu Rev Phytopathol 49, 175–195.

Tan, C.M., Li, C.H., Tsao, N.W., Su, L.W., Lu, Y.T., Chang, S.H., Lin, Y.Y., Liou, J.C., Hsieh, L.C., Yu, J.Z., et al. (2016). Phytoplasma SAP11 alters 3-isobutyl-2-methoxypyrazine biosynthesis in *Nicotiana benthamiana* by suppressing NbOMT1. J Exp Bot 67, 4415–4425.

Taylor, J.S., and Raes, J. (2004). Duplication and divergence: the evolution of new genes and old ideas. Annu Rev Genet 38, 615–643.

Tzfira, T., Vaidya, M., and Citovsky, V. (2004). Involvement of targeted proteolysis in plant genetic transformation by Agrobacterium. Nature 431, 87–92.

Ustun, S., Bartetzko, V., and Bornke, F. (2013). The *Xanthomonas campestris* type III effector XopJ targets the host cell proteasome to suppress salicylic-acid mediated plant defence. PLoS Pathog 9, e1003427.

van Nocker, S., Sadis, S., Rubin, D.M., Glickman, M., Fu, H., Coux, O., Wefes, I., Finley, D., and Vierstra, R.D. (1996). The multiubiquitin-chain-binding protein Mcb1 is a component of the 26S proteasome in *Saccharomyces cerevisiae* and plays a nonessential, substrate-specific role in protein turnover. Mol Cell Biol 16, 6020–6028.

Wang, J.W., Czech, B., and Weigel, D. (2009). miR156-regulated SPL transcription factors define an endogenous flowering pathway in *Arabidopsis thaliana*. Cell 138, 738–749.

Wang, N., Yang, H., Yin, Z., Liu, W., Sun, L., and Wu, Y. (2018). Phytoplasma effector SWP1 induces witches’ broom symptom by destabilizing the TCP transcription factor BRANCHED1. Mol Plant Pathol 19, 2623–2634.

Weintraub, P.G., and Beanland, L. (2006). Insect vectors of phytoplasmas. Annu Rev Entomol 51, 91–111.

Xu, M., Hu, T., Zhao, J., Park, M.Y., Earley, K.W., Wu, G., Yang, L., and Poethig, R.S. (2016). Developmental Functions of miR156-Regulated SQUAMOSA PROMOTER BINDING PROTEIN-LIKE (SPL) Genes in *Arabidopsis thaliana*. PLoS Genet 12, e1006263.

Yoo, S.D., Cho, Y.H., and Sheen, J. (2007). Arabidopsis mesophyll protoplasts: a versatile cell system for transient gene expression analysis. Nature protocols 2, 1565–1572.

Zhang, J., Hogenhout, S.A., Nault, L.R., Hoy, C.W., and Miller, S.A. (2004). Molecular and symptom analyses of phytoplasma strains from lettuce reveal a diverse population. Phytopathology 94, 842–849.

Zhang, X., Zhou, Y., Ding, L., Wu, Z., Liu, R., and Meyerowitz, E.M. (2013). Transcription repressor HANABA TARANU controls flower development by integrating the actions of multiple hormones, floral organ specification genes, and GATA3 family genes in Arabidopsis. Plant Cell 25, 83–101.

